# Gravitational and mechanical forces drive mitochondrial translation

**DOI:** 10.1101/2023.01.18.524628

**Authors:** Taisei Wakigawa, Yusuke Kimura, Mari Mito, Toshiya Tsubaki, Muhoon Lee, Koki Nakamura, Abdul Haseeb Khan, Hironori Saito, Tohru Yamamori, Tomokazu Yamazaki, Akira Higashibata, Tatsuhisa Tsuboi, Yusuke Hirabayashi, Nono Takeuchi-Tomita, Taku Saito, Atsushi Higashitani, Yuichi Shichino, Shintaro Iwasaki

## Abstract

Life on Earth has evolved in a form suitable for the gravitational force of 1 × *g*. Although the pivotal role of gravity in gene expression has been revealed by multiomics approaches in space-flown samples and astronauts, the molecular details of how mammalian cells harness gravity have remained unclear. Here, we showed that mitochondria utilize gravity to activate protein synthesis within the organelle. Genome-wide ribosome profiling revealed reduced mitochondrial translation in mammalian cells and *Caenorhabditis elegans* under both microgravity at the International Space Station and simulated microgravity in a 3D-clinostat on the ground. We found that attenuation of cell adhesion through laminin–integrin interactions causes the phenotype. The downstream signaling pathway including FAK, RAC1, PAK1, BAD, and Bcl-2 family proteins in the cytosol, and mitochondrial fatty acid synthesis (mtFAS) pathway in the matrix maintain mitochondrial translation at high level. Mechanistically, a decreased level of mitochondrial malonyl-CoA, which is consumed by activated mtFAS, leads to a reduction in the malonylation of the translational machinery and an increase in the initiation and elongation of *in organello* translation. Consistent with the role of integrin as a mechanosensor, we observed a decrease in mitochondrial translation via the minimization of mechanical stress in mouse skeletal muscle. Our work provides mechanistic insights into how cells convert gravitational and mechanical forces into translation in an energy-producing organelle.

## Introduction

Since the 1950s, when the first dog was sent to space, biological and medical research has revealed unique responses to life in the space environment ^1^. In addition to radiation, microgravity is one of the main hazards to cells in space and potentially causes oxidative stress ^2,3^, mitochondrial dysfunction ^3–5^, and epigenetic changes ^5,6^. However, due to the limitations of spaceflight experiments, the global view of the cell response to microgravity is still incomplete.

To fill this gap, multi-omics approaches have been applied to a cohort of flight samples, including those from astronauts ^7,8^. These studies explored the epigenome, transcriptome, proteome, and metabolome ^3,9–18^ and revealed microgravity-mediated gene expression profile changes associated with the hazardous phenotypes mentioned above. However, these conventional spaceflight studies have lacked investigations of translational regulation, an acute and inducible means of reprogramming protein expression ^19–22^. Given that translational control is typically followed by transcriptome changes, dynamic shifts in the protein synthesis rate under microgravity could be envisioned, but the “translatome” has not been assessed in previous spaceflight studies.

Here, we comprehensively surveyed the translational response of human tissue cultures and *Caenorhabditis elegans* to microgravity and found that mitochondrial translation is dramatically reduced by microgravity. By elucidating the mechanism of this response, we found that cell adhesion mediated by laminin–integrin and downstream signaling transmission through FAK, RAC1, PAK1, BAD, and Bcl-2 family proteins relays gravitational forces for mitochondrial translation. This leads to activation of the mitochondrial fatty acid synthesis pathway (mtFAS) in the matrix and reduces malonyl-coenzyme A (malonyl-CoA), maintaining a low level of malonylated lysines in translation machinery, which potentiates translation initiation and elongation of mitochondrial protein synthesis. Considering that laminin–integrin functions physiologically as a mechanosensor, the microgravity response may be the manifestation of an impairment of mechanical stress. Consistent with this scenario, we recapitulated the attenuation of mitochondrial translation in unloaded and immobilized mouse hindlimbs. This study revealed the previously uncharacterized unique connection between the extracellular environment and mitochondrial translation.

## Results

### Mitochondrial translation repression under microgravity in space

To investigate the translational landscape under microgravity, we conducted genome-wide ribosome profiling ^23,24^ on HEK293 cells cultured at the International Space Station (ISS) (Figure 1A). For this purpose, cells were first cultured at 1 × *g,* artificially generated by centrifugation in the ISS, and then under microgravity conditions for 24 and 48 h (Figure 1A). In contrast, control cells were maintained at 1 × *g* until harvesting. The cells were treated with translation inhibitors (cycloheximide and chloramphenicol), frozen, and returned to the laboratory on Earth for library preparation.

**Figure 1.**
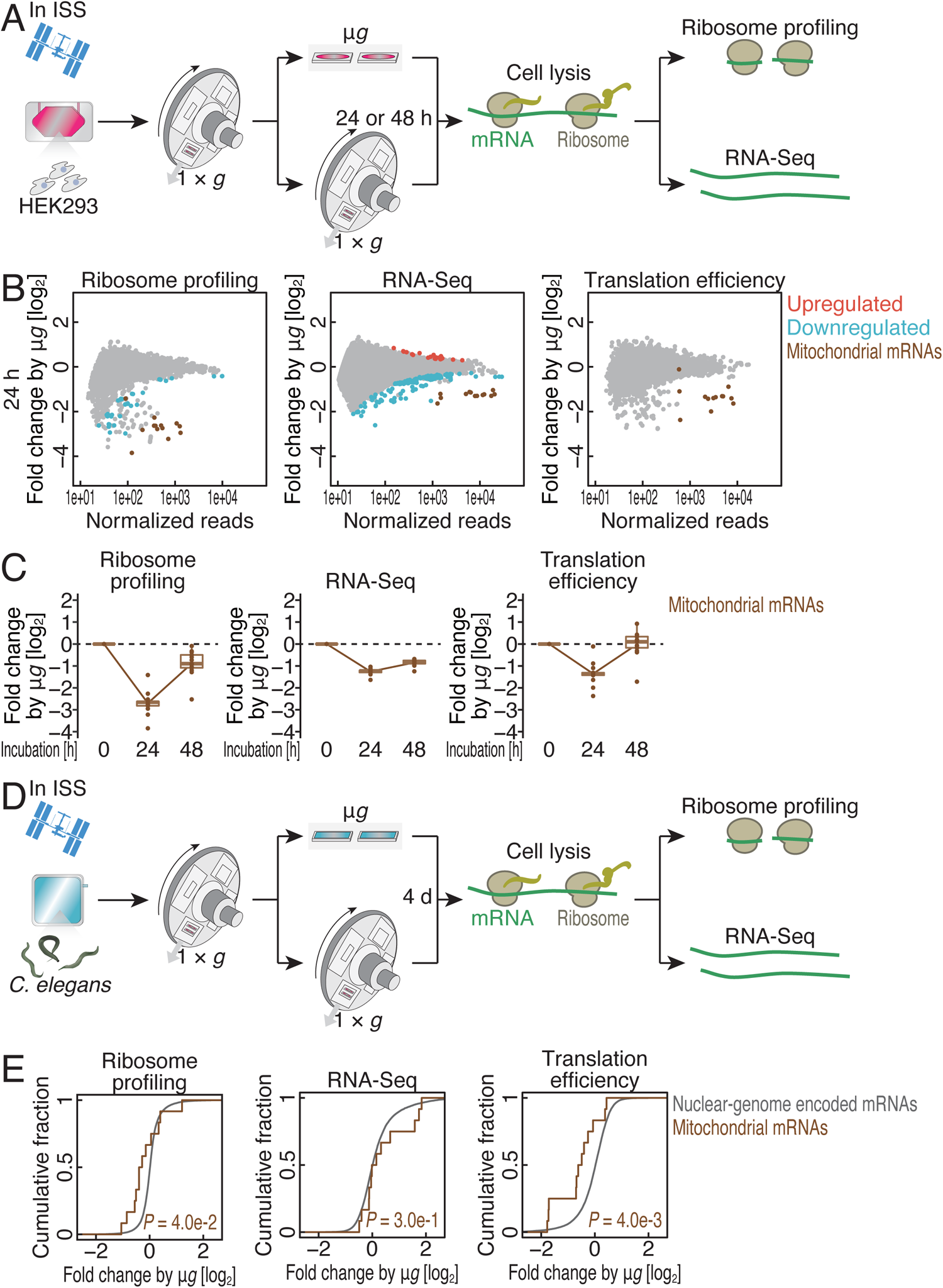
Genome-wide ribosome profiling reveals mitochondrial translation repression by microgravity in humans and nematodes. (A and D) Schematic representation of the experiments in the international space station (ISS) for HEK293 cells (A) and *C. elegans* (D). µ*g*: microgravity. (B) MA (M, log ratio; A, mean average) plots for ribosome footprint change (left), RNA abundance change (middle), and translation efficiency change (right) after 24-h microgravity (μ*g*) culture in HEK293 cells. Significantly altered transcripts (false discovery rate [FDR]< 0.05) and mitochondrial genome-encoded mRNAs are highlighted. (C) Box plots for ribosome footprint change (left), RNA abundance change (middle), and translation efficiency change (right) in mitochondrial genome-encoded mRNAs over the course of microgravity culture. (E) Cumulative distributions of ribosome footprint change (left), RNA abundance change (middle), and translation efficiency change (right) in *C. elegans* with respect to nuclear genome-encoded mRNAs and mitochondrial genome-encoded mRNAs after 4-d microgravity culture. The p value was calculated by the Mann‒Whitney *U* test. In box plots (C), the median (centerline), upper/lower quartiles (box limits), and 1.5× interquartile range (whiskers) are shown. See also Figure S1 and Table S2.

Due to spaceflight restrictions, the amount of material available was limited for standard ribosome profiling experiments. To overcome this issue, we harnessed Thor-Ribo-Seq, a ribosome profiling derivative tailored for low input based on RNA-dependent RNA amplification by *in vitro* transcription ^25^. The application allowed us to obtain high-quality data from samples returned from the ISS, which presented hallmarks of ribosome footprints: a sharp peak in the read length at 28-29 nt (Figure S1A), 3-nt periodicity (Figure S1B), and high reproducibility (Figure S1C left).

Our analysis revealed the impacts of microgravity on translation, especially a reduction in a subset of mRNAs (Figures 1B left, S1D left, and S1E). Among those, we found that protein synthesis from mitochondrial mRNAs, which are encoded in the organelle genome, was remarkably sensitive to microgravity (Figure 1B left). Since our method captured footprints from both cytosolic ribosomes (cytoribosomes) and mitochondrial ribosomes (mitoribosomes) ^26,27^, we could interrogate both translation systems simultaneously. The reduction in mitoribosome footprints could not be explained by a reduction in the corresponding mRNAs, as measured by RNA sequencing (RNA-Seq) of the same materials (Figures 1B middle, S1C right, S1D middle, and S1E). Thus, the translation efficiency, calculated by over- or underrepresentation of ribosome footprints over RNA-Seq fragments, still showed a decrease in mitochondrial mRNAs (Figure 1B right). In contrast, the translation of mitochondrial proteins encoded in the nuclear genome was insensitive to microgravity (Figure S1F-G).

Mitochondrial translation regulation may be an early response to microgravity. We observed that the change in mitochondrial translation was more dramatic after 24 h of incubation compared to 48 h under microgravity conditions (Figure 1C) (see Discussion for details).

To further extend our exploration from cell culture to the whole-body level, we used nematode samples flown in the ISS ^16^. The L1 larvae were cultured under microgravity conditions for 4 d, while the control worms were incubated in a 1 × *g* centrifuge (Figure 1D). Similar to human cells, ribosome profiling and RNA-Seq in nematodes (Figure S1I-K) revealed translational alterations in a subset of mRNAs (Figure S1L-M). Remarkably, we observed a reduction in mitochondrial translation efficiency (Figure 1E), as shown by the cell-based experiments.

Through these spaceflight experiments, we concluded that attenuation of mitochondrial translation is a pervasive response to microgravity in higher eukaryotes.

### The mitochondrial translation response is recapitulated by simulated microgravity on the ground

To survey microgravity-mediated mitochondrial translation repression in the laboratory on the ground, we harnessed a clinostat that rotates the cell culture flask in three dimensions to reduce gravitational forces on the cells and simulate microgravity conditions (Figure 2A). Ribosome profiling and RNA-Seq of HEK293 cells revealed that simulated microgravity phenocopied the attenuated mitochondrial translation observed in the ISS (Figures 2B-C and S2A); we detected the reduction in mitoribosome footprints even after 1 h of simulated microgravity treatment (Figure 2C). Again, we did not find a significant impact on nuclear genome-encoded mitochondrial proteins (Figure S2B and S2C). Gravity-mediated mitochondrial translation regulation was further ensured by mitochondria-specific fluorescent noncanonical amino acid tagging on gels (on-gel mito-FUNCAT) ^28–32^ (Figure 2D). Since the same molecular phenotype was observed in human cells (HEK293) and mouse cells (C2C12 and 3T3) (Figures 2D and S2D), the mitochondrial translation response to microgravity is likely driven by conserved mechanisms.

**Figure 2.**
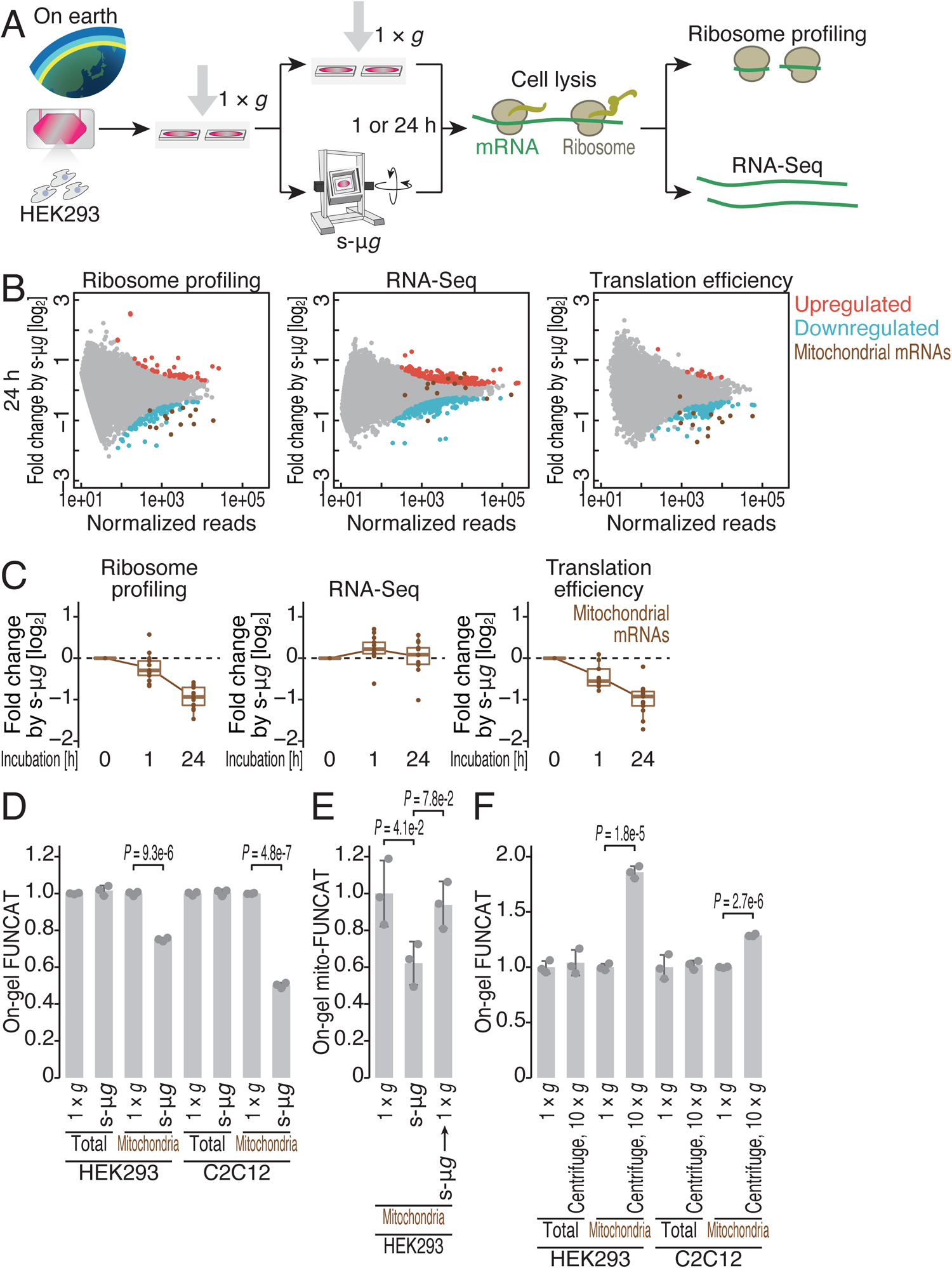
Mitochondrial translation is a function of gravity. (A) Schematic representation of experiments with a 3D clinostat to induce simulated microgravity (s-μ*g*) in HEK293 cells. (B) MA plots for ribosome footprint change (left), RNA abundance change (middle), and translation efficiency change (right) after 24-h simulated microgravity culture in HEK293 cells. Significantly altered transcripts (false discovery rate [FDR] < 0.05) and mitochondrial genome-encoded mRNAs are highlighted. (C) Box plots for ribosome footprint change (left), RNA abundance change (middle), and translation efficiency change (right) in mitochondrial genome-encoded mRNAs over the course of simulated microgravity culture. (D) On-gel FUNCAT experiments to monitor total and mitochondrial translation in the indicated cell lines and conditions. The cells were cultured under s-μ*g* for 24 h before cell harvesting. (E) On-gel FUNCAT experiments examining mitochondrial translation in the indicated cell lines and conditions. For the recovery experiments, the cells were incubated under s-μ*g* for 24 h and then at 1 × *g* for 24 h. (F) On-gel FUNCAT experiments to monitor total and mitochondrial translation in the indicated cell lines and conditions. The cells were cultured at 10 × *g* for 24 h before cell harvesting. For D-F, the data from three replicates (points), the mean values (bars), and the s.d.s (errors) are shown. The p values were calculated by Student’s t test (two-tailed) (D and F) and by the Tukey‒Kramer test (two-tailed) (E). In box plots (C), the median (centerline), upper/lower quartiles (box limits), and 1.5× interquartile range (whiskers) are shown. See also Figure S2 and Table S2.

Our on-gel mito-FUNCAT experiments showed that mitochondrial translation is a function of gravity. When the cell culture was returned to standard 1 × *g*, the reduction in mitochondrial translation induced by microgravity was restored to the baseline level, suggesting the elasticity of the mechanism (Figure 2E). Moreover, centrifugation-induced 10 × *g* hypergravity resulted in the activation of mitochondrial translation (Figure 2F), whereas centrifugation at 1 × *g* did not (Figure S2E).

Given that microgravity induces mitochondrial stress ^2–4^, we investigated the correspondence between these stresses and *in organello* protein synthesis under microgravity. We did not observe a significant alteration in the copy number of mitochondrial DNA, which is susceptible to oxidative damage, under simulated microgravity (Figure S2F-G). Additionally, our microscopy analysis did not reveal mitochondrial fragmentation, which can be induced by mitochondrial oxidative stress ^33,34^

(Figure S2H-I). In both space and clinostat samples, we could not find a signature of the mitochondrial unfolded protein response (mtUPR) ^35^ since the transcriptomic alteration of targets ^36^ was limited (Figures S1H and S2J). Thus, our data indicated that the repression of mitochondrial translation is not caused by oxidative stress or morphological defects in the organelle.

### The cell adhesion pathway relays gravitic force to mitochondrial translation

How do cells recognize gravity and transmit the information to organelle translation? Recent studies have highlighted the attenuation of cell adhesion and subsequent signal transduction through the focal adhesion kinase (FAK) pathway by microgravity ^37–39^. Consistent with these reports, we observed that the phosphorylation of FAK, which is driven by cell adhesion, was reduced by simulated microgravity (Figure S3A). Thus, we reasoned that modulated cell adhesion may reduce mitochondrial translation in microgravity.

We first investigated the relationship between cell adhesion and organelle translation under regular gravity. Considering the translation flux of the α and β subunits of integrin, a receptor of the extracellular matrix (ECM), the laminin receptor α_6_β_1_ was an abundant integrin class in HEK293 cells (Figure S3B). Accordingly, pretreatment of cell culture dishes with laminin-511 (or α_5_β_1_γ_1_), which has a high affinity for integrin α_6_β_1_ ^40,41^, enhanced cell adhesion in a dose-dependent manner (Figure 3A and 3B top) and thus elevated FAK phosphorylation (Figure S3C). Through on-gel mito-FUNCAT, we found that mitochondrial protein synthesis was concomitantly activated by increasing the laminin concentration (Figure 3A and 3B bottom left) and correlated with cell adhesion activity (Figure 3B bottom right). Similar mitochondrial translation activation by laminin was also observed in other cell types (human HAP1 and mouse C2C12 and 3T3) (Figures 3C, 4I, and S3D). Conversely, mitochondrial translation was inhibited by the RGDS peptide, an integrin inhibitor (Figure 3D), which led to FAK dephosphorylation (Figure S3E). In contrast to *in organello* translation, total protein synthesis, which predominantly originates from cytosolic translation, was not altered by laminin or RGDS peptide treatment (Figure 3B-D). These results showed that cell adhesion specifically enhances mitochondrial translation.

**Figure 3.**
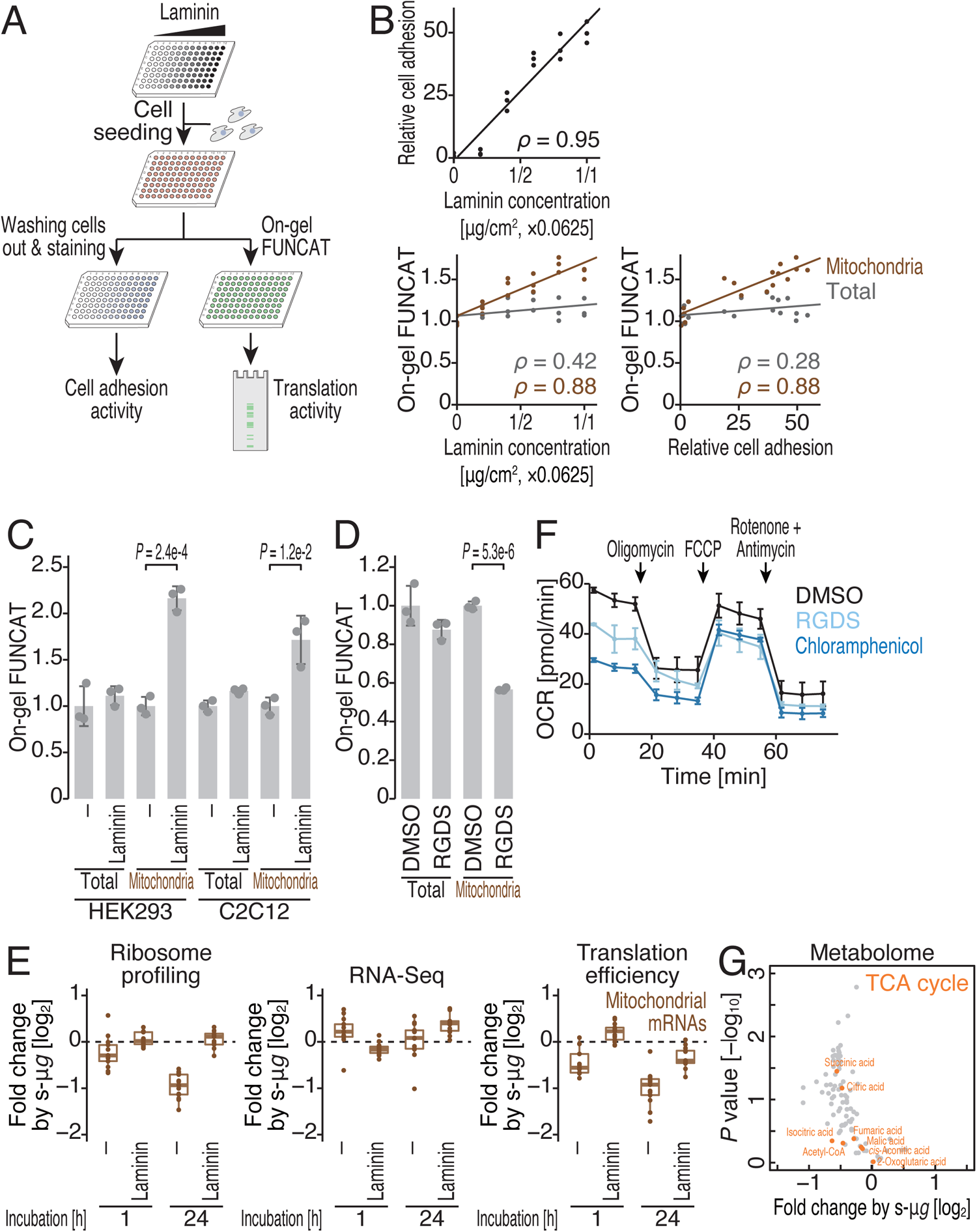
The attenuation of the laminin–integrin signaling pathway represses mitochondrial translation under microgravity. (A) Schematic representation of the experiments to investigate the impacts of laminin– integrin-mediated cell adhesion on mitochondrial translation. (B) Relationships among laminin concentration, cell adhesion activity, and translation activity. For cell adhesion activity, the mean value without laminin precoating was set to “1”. Total and mitochondrial protein synthesis were separately assayed. ρ, Spearman’s rank correlation coefficient. (C) On-gel FUNCAT experiments to monitor total and mitochondrial translation in the indicated cell lines and conditions. (D) On-gel FUNCAT experiments to monitor total and mitochondrial translation in the indicated cell lines and conditions. The cells were treated with 10 μg/ml RGDS peptide for 24 h. (E) Box plots for ribosome footprint change (left), RNA abundance change (middle), and translation efficiency change (right) in mitochondrial genome-encoded mRNAs over the course of simulated microgravity culture with or without laminin precoating. Note that the data without laminin precoating are the same as those in Figure 2C. (F) Oxygen consumption rate (OCR) of HEK293 cells at basal respiration and after treatment with oligomycin, carbonyl cyanide 4-(trifluoromethoxy)phenylhydrazone (FCCP), or rotenone/antimycin. (G) Volcano plot for the metabolomic change after 24-h simulated microgravity culture. TCA cycle-associated metabolites are highlighted. For C and D, the data from three replicates (points), the mean values (bars), and the s.d.s (errors) are shown. The p values were calculated by Student’s t test (two-tailed). For F, the mean values (points) from three replicates and the s.d.s (errors) are shown. In box plots (E), the median (centerline), upper/lower quartiles (box limits), and 1.5× interquartile range (whiskers) are shown. The experiments were conducted in HEK293 cells unless otherwise noted. See also Figure S3 and Table S2.

**Figure 4.**
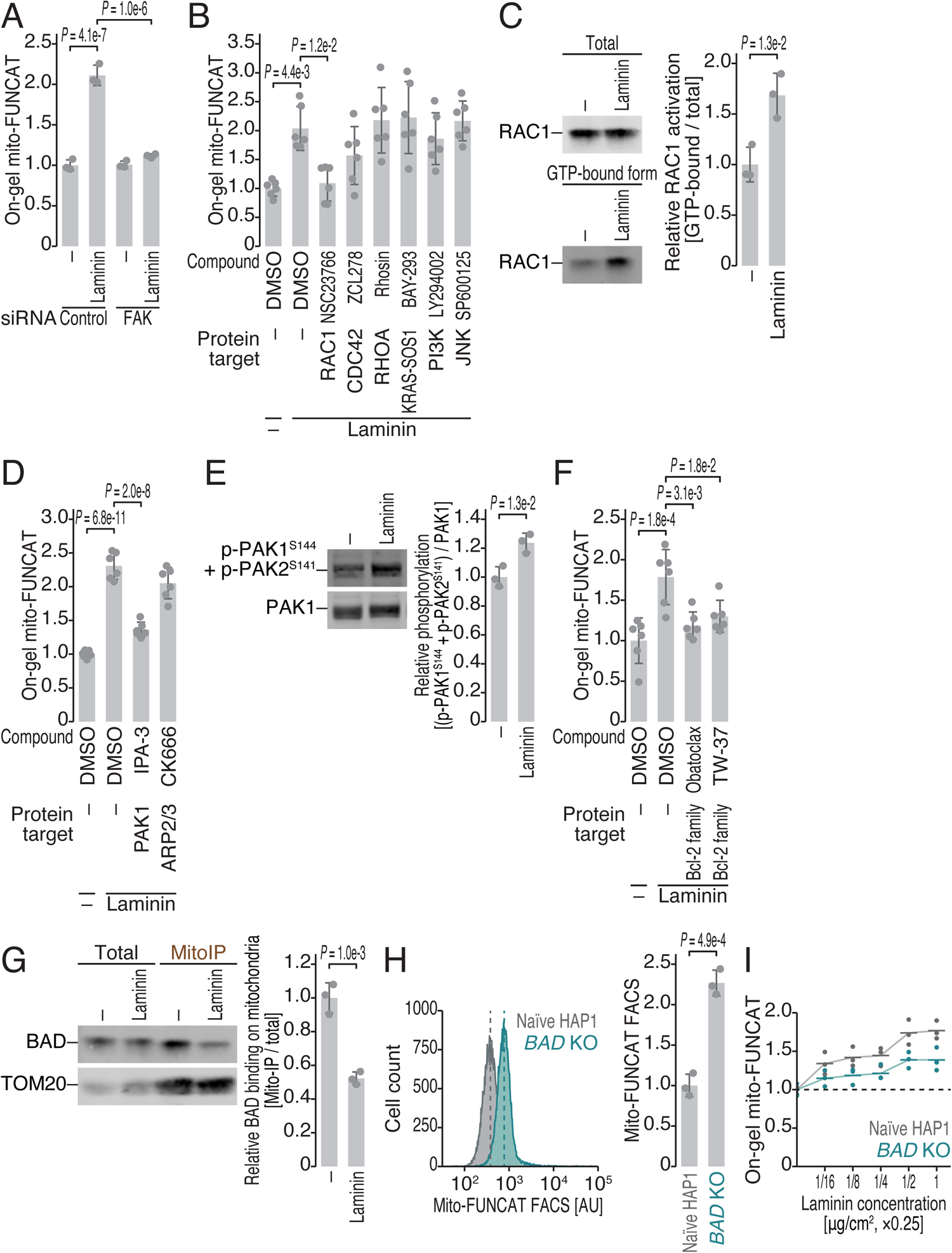
The cellular pathway that transduces the signal from laminin–integrin cell adhesion to mitochondrial translation activation. (A) On-gel FUNCAT experiments to monitor mitochondrial translation after siRNA transfection to knock down FAK. (B, D, and F) On-gel FUNCAT experiments to monitor mitochondrial translation under treatment with the indicated compounds at the following concentrations: NSC23766, 200 μM; ZCL278, 100 μM; Rhosin, 50 μM; BAY-293, 1 μM; LY294002, 10 μM; SP600125, 20 μM; IPA-3, 5 μM; CK666, 25 μM; obatoclax, 2 μM; and TW-37, 10 μM. (C) Western blotting of RAC1 protein in the total cell lysate and in the GTP-bound fraction pulled down with the p21-binding domain (PBD) of PAK1. The quantified relative amount of RAC1 in the GTP-bound, activated form is shown. (E) Western blotting of total PAK1 protein and the phosphorylated form (at S144 and S141). The quantified relative amount of phosphorylated PAK1 is shown. (G) Western blotting of BAD protein in the total lysate and in the mitochondrial immunoprecipitation (MitoIP) fraction with an anti-TOM22 antibody. (H) Representative distribution of Cy3-conjugated HPG signals normalized to AF647-labeled TOMM20 signals in the indicated cell lines (left). The dashed line represents the mean of the distribution. The quantification is shown on the right. The distribution of unnormalized Cy3-conjugated HPG signals and AF647-labeled TOMM20 signals are shown in Figure S4J. (I) On-gel FUNCAT experiments to monitor mitochondrial translation in naïve and *BAD* KO HAP1 cells with a titrated concentration of laminin for precoating. For A-H, the data from replicates (points, n = 3 for A, C, E, G, and H; n = 6 for B, D, and F), the mean values (bars), and the s.d.s (errors) are shown. The p values were calculated by Student’s t test (two-tailed) (C, E, G, and H) and by the Tukey‒Kramer test (two-tailed) (A, B, D, and F). For I, the data from three replicates and the mean values (bars) are shown. The experiments were conducted in HEK293 cells unless otherwise noted. See also Figure S4.

To test whether microgravity-mediated repression of mitochondrial translation is caused by reduced cell adhesion, we pretreated a cell culture flask with laminin, cultivated the cells under simulated microgravity, and then conducted ribosome profiling and RNA-Seq. Strikingly, laminin counteracted the effect of microgravity on mitochondrial protein synthesis (Figure 3E). We concluded that cell adhesion is a mediator of gravitational forces for mitochondrial translation.

The loss of *in organello* translation homeostasis results in physiological dysfunction of the organelle. Attenuation of cell adhesion by RGDS peptides hampered the oxygen consumption rate of the mitochondria (Figure 3F), as did the mitochondrial translation inhibitor chloramphenicol. Moreover, metabolites associated with the tricarboxylic acid (TCA) cycle were reduced by simulated microgravity (Figure 3G). These data suggested that gravity is required to maintain the metabolic integrity of the organelle.

### The FAK–RAC1–PAK1–BAD–Bcl-2 family protein axis conveys a signal from cell adhesion to mitochondrial protein synthesis

Given the phosphorylation and activation of FAK upon laminin–integrin interaction ^42,43^ (Figure S3C), we investigated the significance of this factor for mitochondrial translation. Indeed, FAK knockdown resulted in the loss of laminin-mediated activation of *in organello* translation (Figures 4A and S4A).

We next asked how the downstream signaling pathway(s) of FAK connects to mitochondria. Since FAK influences widespread enzymes such as small GTPases (RAC1, CDC42, RHOA, and KRAS), phosphatidylinositol 3-kinase (PI3K), and Jun N-terminal kinase (JNK) ^42,43^ (Figure S4B), these activities were pharmacologically inhibited (Figure S4C), and the potential for laminin-mediated mitochondrial translation activation was evaluated via on-gel mito-FUNCAT. Screening showed that RAC1 inhibition specifically hampered laminin-mediated mitochondrial translation activation (Figures 4B and S4D). As expected, laminin treatment led to the accumulation of the active GTP-bound form of RAC1 (Figure 4C), which is evoked by phosphorylated FAK ^42,43^.

Further compound treatment narrowed the downstream pathway of RAC1 ^44^ to PAK1 but not to ARP2/3 (Figures 4D and S4E-F). We observed elevated phosphorylation of PAK1, which is mediated by GTP-activated RAC1 ^44^, in laminin-treated cells (Figure 4E).

Given that PAK1 couples with the pro-survival cascade driven by Bcl-2 family proteins on the mitochondrial outer membrane ^45–47^, we reasoned that the Bcl-2 family pathway may participate in cell adhesion-mediated mitochondrial translation activation. We treated cells with pan-Bcl-2 family inhibitors (such as obatoclax and TW-37) and found that the effect of laminin on mitochondrial protein synthesis was counteracted (Figures 4F and S4G-H).

To regulate Bcl-2 family proteins, PAK1 phosphorylates BAD ^48^, one of the pro-apoptotic Bcl-2 homology 3 (BH3)-only proteins that bind to Bcl-2 family proteins on the mitochondrial outer membrane and inhibits the pro-survival effect ^49–51^. BAD phosphorylation reduces its affinity for Bcl-2 family proteins, dissociating BAD from Bcl2 proteins and thus from the mitochondrial outer membrane ^49–51^. Indeed, laminin treatment decreased the level of the mitochondria-associated fraction of BAD (Figure 4G).

Therefore, we examined the role of BAD in mitochondrial translation. During our analysis of the knockout of *BAD* in HAP1 cells (Figure S4I), we noticed a reduction in mitochondrial mass in this cell line (Figure S4J bottom). To normalize the mitochondrial volume and simultaneously determine the extent of protein synthesis in the organelle, we harnessed mito-FUNCAT coupled with fluorescence-activated cell sorting (FACS) ^31^. The mito-FUNCAT FACS revealed that the deletion of *BAD* resulted in a high level of net mitochondrial translation due to the loss of the suppressive mechanism for mitochondrial translation (Figures 4H and S4J). Ultimately, *BAD* deficiency led to an attenuated response to laminin in mitochondrial translation (Figure 4I).

Taken together, our results revealed the signal transduction pathway from the extracellular matrix laminin to the Bcl-2 family protein on the mitochondrial outer membrane for *in organnello* translation activation.

### Laminin-mediated mitochondrial translation activation is distinct from known mechanisms

These findings prompted us to investigate the key molecular events in the matrix where mitochondrial translation occurs. For this purpose, we tested the possibility of the involvement of known regulatory mechanisms of mitochondrial translation.

The membrane potential is a prerequisite for mitochondrial protein synthesis ^52–54^ (Figure S5A). However, alterations in the membrane potential could not explain the dynamics of mitochondrial translation upon cell adhesion modulation (Figure S5B-E).

Mitochondrial translation has been reported to occur at cristae membranes ^30^, where OXPHOS complexes are enriched ^55–57^. Indeed, we observed that more cristae formed with laminin treatment through electron microscopy (Figure S5F-H). Moreover, the loss of the inner membrane-located OPA1, which leads to oligomerization and subsequent tubulation of the inner membrane to form cristae structure ^58^, in MEF cells ^59^ reduced the basal level of mitochondrial translation (Figure S5I). However, the *OPA1* KO cells still retained the potential to activate mitochondrial translation upon laminin treatment (Figure S5I-J), indicating that the two translation regulatory mechanisms are independent. Thus, we concluded that the increase in cristae upon laminin treatment was a result (not a cause) of activated mitochondrial translation, probably due to increased formation of respiratory chain super complexand subsequent inner-membrane curvature and tubulation ^60^.

In yeast, rapid communication between the cytosolic and mitochondrial translation systems has been reported ^61^; cytosolic translation synchronizes its output with mitochondrial protein synthesis to maintain the synthesis rate of OXPHOS subunits. However, our ribosome profiling did not detect a large alteration in OXPHOS complex subunits encoded in the nuclear genome (Figures S1F and S2B). Moreover, a recent report suggested that the synchronization mechanism is not conserved in humans ^62^.

### mtFAS-coupled mitochondrial translation activation

We subsequently tested whether mitochondrial fatty acid synthesis (mtFAS) (Figure 5A) is involved in laminin-induced mitochondrial translation. mtFAS exploits manifold enzymes to covalently extend acyl chains on acyl carrier proteins (ACPs) (Figure 5A) ^63^. The eight-carbon fatty acid octanoate on ACP is converted into lipoic acid, generating lipoylated proteins (Figure 5A) ^63^. Since the loss of this metabolic pathway suppresses *in organello* translation in yeasts ^63–65^, we reasoned that the involvement of mtFAS in laminin-mediated mitochondrial translation activation. Indeed, we found that laminin treatment increased mtFAS activity, assessed by lipoylation (Figure 5B). Conversely, blocking the downstream signal transduction pathway with the Bcl-2 family inhibitor TW-37 was associated with low mtFAS efficiency (Figure 5C).

**Figure 5.**
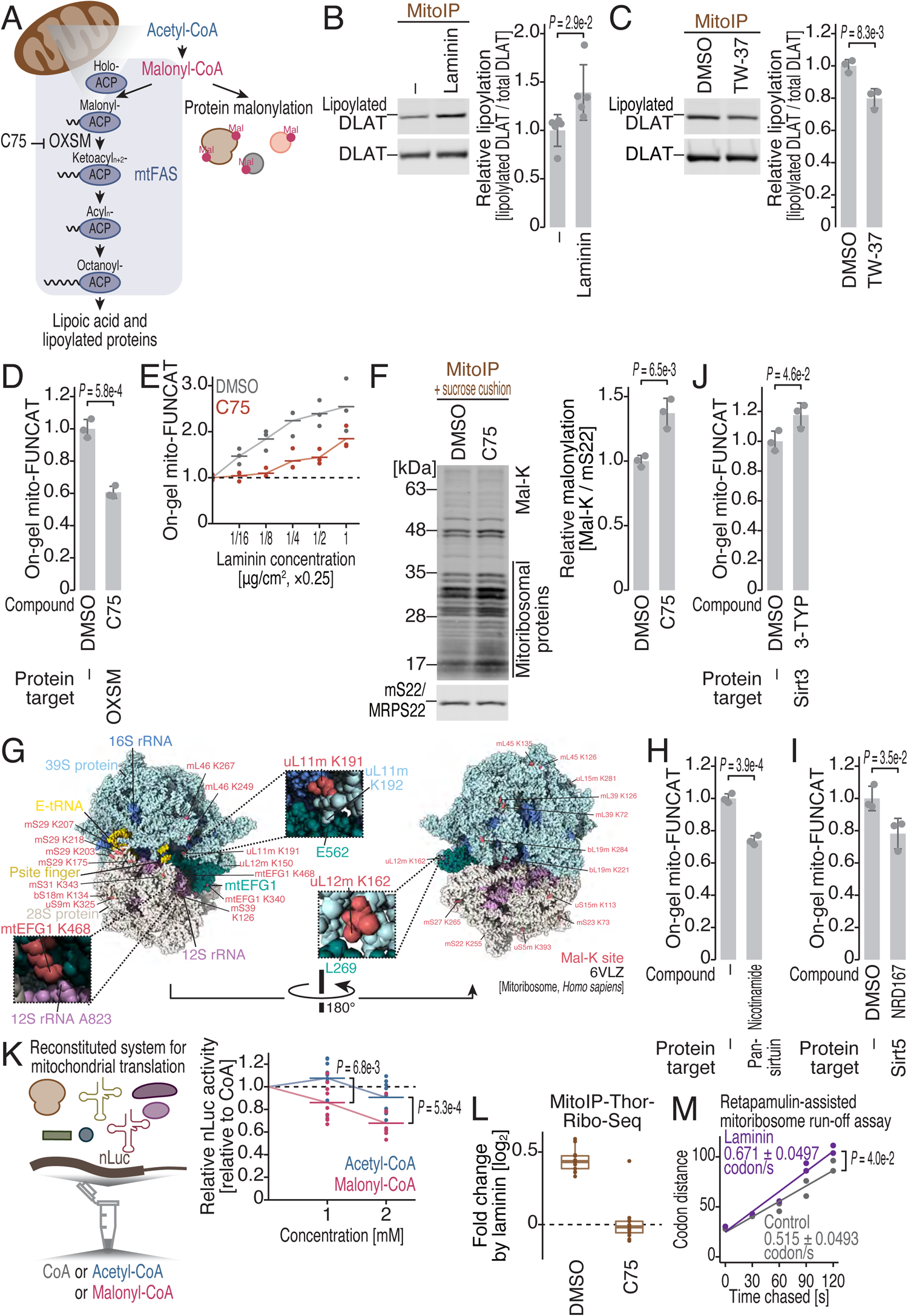
mtFAS balances manlonyl-CoA for malonylation of the mitochondrial translation machinery and thus for translation *in organello*. (A) Schematic representation of acetyl-CoA, malonyl-CoA, and the downstream pathways in the mitochondrial matrix. (B and C) Western blotting of DLAT and lipoylated DLAT proteins under the indicated conditions. The quantified relative amount of lipoylated DLAT is shown. (D and H-J) On-gel FUNCAT experiments to monitor mitochondrial translation under treatment with the indicated compounds at the following concentrations: C75, 50 μM; nicotinamide, 10 mM; NRD167, 10 μM; and 3-TYP, 10 µM. (E) On-gel FUNCAT experiments to monitor mitochondrial translation in C75-treated cells with a titrated concentration of laminin for precoating. (F) Western blotting of proteins with malonylated lysine (Mal-K) in the mitochondrial IP (MitoIP) fraction with an anti-TOM22 antibody and then the pellet of the sucrose cushion ultracentrifugation fraction. MRPS22 (mS22) was used as the loading control. The quantified relative amount of Mal-K signal is shown. (G) Malonylated lysine residues (Mal-K) found in the compendium of protein lysine modifications (CPLM) database ^68^ were mapped on the mitoribosome complexed with mtEFG1 (6VLZ) ^88^, using UCSF ChimeraX ^106^. (K) *In vitro* translation of mammalian mitochondria reconstituted with recombinant factors. CoA, acetyl-CoA, or malonyl-CoA was added to the reaction mixture. A representative nLuc reporter mRNA was used to probe basal translation activity in the system. Figure S6L shows the effect of CoA on the reaction. (L) High-resolution measurement of mitochondrial translation with MitoIP-Thor-Ribo-Seq. The fold change in mitoribosome footprints by laminin precoating was measured in the presence or absence of C75. (M) Mitochondrial translation elongation rate measured by the mitoribosome run-off assay with retapamulin and MitoIP-Thor-Ribo-Seq. The regression line for the expansion of the mitoribosome-free area over time determines the mitochondrial translation elongation rates in the presence or absence of laminin precoating. For B-D, F, and H-J, the data from replicates (points, n = 5 for B; n = 3 for C, D, and H-J), the mean values (bars), and the s.d.s (errors) are shown. The p values were calculated by Student’s t test (two-tailed). For E and K, the data from three replicates (E) and nine replicates (K) (points) and the mean values (bars) are shown. The p values in K were calculated by the Tukey‒Kramer test (two-tailed). For M, the p value was calculated by two-way ANOVA. In box plots (L), the median (centerline), upper/lower quartiles (box limits), and 1.5× interquartile range (whiskers) are shown. All the experiments except for K were conducted in HEK293 cells. See also Figure S5-6 and Table S1-2.

Harnessing C75, an inhibitor of the 3-ketoacyl-ACP synthase OXSM (Figure 5A) ^66^, we found that the inactivation of mtFAS (Figure S6A) led to reduced basal mitochondrial translation (Figure 5D). Strikingly, C75 treatment attenuated the translational response to laminin (Figure 5E). Thus, mtFAS activity is coupled with mitochondrial translation.

### Protein demalonylation in mitochondria leads to mitochondrial translation

Despite the reported linkage between mtFAS and mitochondrial translation in yeasts ^63^, the underlying mechanism has remained largely unknown. In the cytosol, the counterpart fatty acid synthase (FASN) balances the abundance of malonyl-CoA; otherwise, excess malonyl-CoA generated by FASN impairment induces malonylation on lysine residues ^67^. Given that, we envisioned a similar scenario in mitochondria (Figure 5A). We observed that laminin-mediated mtFAS activation (Figure 5B) is associated with the demalonylation of mitochondrial proteins, especially mitochondrial translational machinery (such as mitoribosomal proteins) (Figure S6B, MitoIP + sucrose cushion). Conversely, mtFAS inhibition by C75 increased the malonylated fraction of the translational machinery proteins (Figure 5F). Through the lysine modifications assessed by mass spectrometry ^68^, diverse solvent-accessible lysines in mitochondrial ribosome proteins, elongation factors, and recycling factors have been identified as malonylation sites (Figures 5G and S6C-E, and Table S1), suggesting their functional implications in translation regulation.

To test the importance of mitochondrial protein malonylation in the regulation of protein synthesis, we directly modulated the protein malonylation status in mitochondria. Sirt5, a sirtuin family protein, is a major demalonylase in mitochondria ^69–71^. Chemical perturbation of this protein by the pan-sirtuin inhibitor nicotinamide or by the Sirt5-specific inhibitor NRD167 suppressed mitochondrial translation (Figure 5H-I), leading to hyper-malonylation of the translation machinery (Figure S6F) independent of mtFAS alteration (Figure S6G-H). Notably, in combination with laminin treatment, Sirt5 inhibition had limited effects on mitochondrial translation (Figure S6I-J), probably due to the unavailability of malonyl-CoA for further hyper-malonylation by activated mtFAS. In addition to malonylation, lysines are often subjected to acetylation by acetyl-CoA donors in mitochondria ^71^. However, the inhibition of Sirt3, a mitochondrial deacetylase ^71–73^, by 3-TYP did not hamper mitochondrial translation but rather increased mitochondrial translation (Figures 5J and S6K), suggesting the acyl group-dependent function of the lysine modifications for i*n organello* translation.

A high concentration of acetyl-CoA/malonyl-CoA in the matrix leads to nonenzymatic lysine acylation ^71,74^. To investigate the direct effect of acetyl-CoA and malonyl-CoA on mitochondrial protein synthesis, we employed a reconstituted translation system with recombinant factors (Figure 5K left) ^75,76^. We observed that, compared to the CoA control, malonyl-CoA inhibited mitochondrial translation (Figures 5K right and S6L), whereas acetyl-CoA had a relatively weaker impact. Taken together, these data led us to conclude that a low malonyl-CoA supply resulting from activated mtFAS and the subsequent reduction in lysine malonylations in the translation machinery leads to enhanced mitochondrial protein synthesis.

### Both initiation and elongation are controlled by protein malonylation

Given that malonylated lysines were found in mitoribosomal proteins close to initiation/elongation factors or those factors *per se* (Figures 5G and S6C-D, and Table S1), we investigated the impact of this regulatory system on mitochondrial translation initiation and elongation. For this purpose, we employed MitoIP-Thor-Ribo-Seq, a high-resolution ribosome profiling derivative tailored for mitochondrial translation ^77^. Since translation initiation is a general rate-limiting step of mitochondrial protein synthesis ^77^, the overall number of mitoribosome footprint reads represents a proxy for initiation speed. We observed a global increase in the mtioribosome loading on mRNAs following laminin treatment (Figure 5L), whereas C75 blocked this phenotype.

We also applied a retapamulin-assisted mitoribosome run-off assay with MitoIP-Thor-Ribo-Seq ^77^ to monitor the elongation rate. Retapamulin traps mitoribosomes only at start codons and allows active elongation of mitoribosomes to complete protein synthesis. Thus, a time-course chase with chloramphenicol after retapamulin treatment generates a mitoribosome-free area downstream of the start codon, measuring the translation elongation rate ^77^. This assay revealed that the elongation of mitochondrial ribosomes was enhanced by laminin treatment (Figure 5M). Thus, both the initiation and elongation steps of mitochondrial translation were regulated through the cell adhesion-mtFAS axis.

### Mechanical stress enhances mitochondrial protein synthesis in vivo

The correspondence between mitochondrial translation and laminin–integrin-mediated cell adhesion led us to study the physiological importance of this cascade. Considering that integrin plays a critical role as a mechanosensor ^78–81^, we hypothesized that mechanical stress promotes mitochondrial translation. Given that mechanical stress and gravity maintain skeletal muscle homeostasis and FAK phosphorylation ^37–39,82,83^, we focused on translation in the tissues of mice. In a minimized mechanical stress (MMS) model, mice underwent hindlimb unloading and immobilization, and the soleus was subjected to ribosome profiling and RNA-Seq (Figure 6A). Along the duration of MMS, we found reduced mitochondrial translation at 14 d after the start of minimized mechanical stress (Figure 6B-C), the time at which we detected reduced muscle weight (Figure S7A). Our results indicated the importance of mechanical stress for mitochondrial protein synthesis.

**Figure 6.**
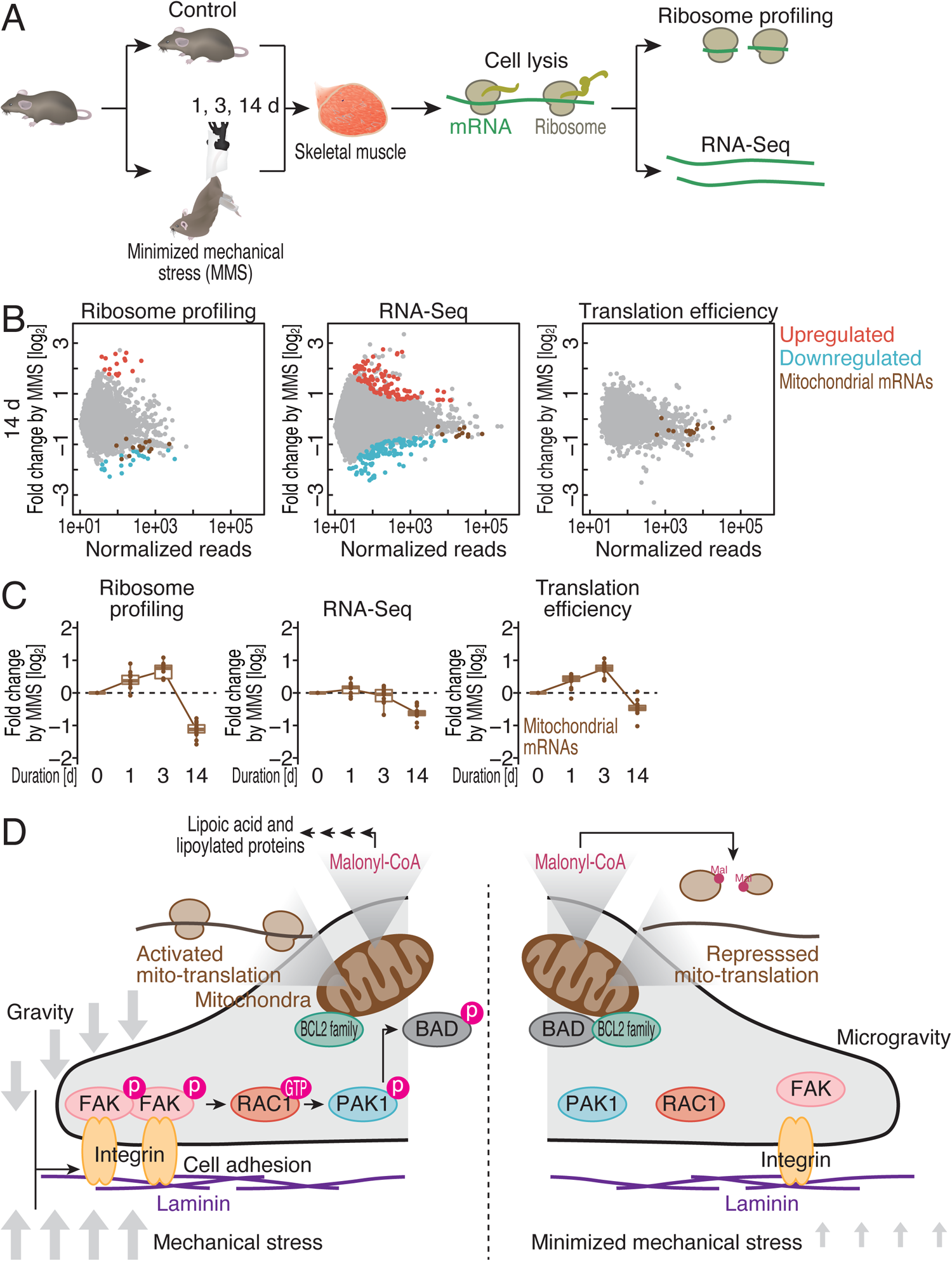
Minimized mechanical stress reduces mitochondrial translation. (A) Schematic representation of experiments with minimized mechanical stress (MMS) in mouse hindlimbs. (B) MA plots for ribosome footprint change (left), RNA abundance change (middle), and translation efficiency change (right) after 14-d MMS in the mouse soleus. Significantly altered transcripts (false discovery rate [FDR] < 0.05) and mitochondrial genome-encoded mRNAs are highlighted. (C) Box plots for ribosome footprint change (left), RNA abundance change (middle), and translation efficiency change (right) in mitochondrial genome-encoded mRNAs over the course of MMS. (D) Schematic representation of the model of gravitational and mechanical force-mediated mitochondrial translation activation. See also Figure S7 and Table S2.

## Discussion

Starting from the translatome analysis of space-flown samples, we elucidated a previously unexplored cell signaling pathway that connects cell adhesion sensing, the mitochondrial malonyl-CoA balance, and *in organello* translation (Figure 6D). Although translational reprogramming mediated by integrins and mechanical forces has been suggested for decades ^84–86^, the landscape has largely not been elucidated. Our study revealed mitochondrial translation as a subject of the regulation. In muscle, which is constantly subjected to mechanical stresses through loading, postural maintenance, and exercise, this system enables rational regulation of energy production in mitochondria: increased mechanical stresses lead to increased mitochondrial protein synthesis and thus modulate the organelle’s functions.

Although the short-term microgravity conditions in our study did not provide strong evidence, long-term culture under microgravity may ultimately lead to mitochondrial damage. Indeed, the recent multiomics approach utilizing data from NASA’s GeneLab platform, which included an astronaut cohort and space-flown samples ^3^, illustrated mitochondrial damage as a hallmark of spaceflight. The attenuation of mitochondrial translation may prime the damage processes in space.

Intriguingly, after 48 h of microgravity exposure, we observed recovery of translation efficiency for mitochondrial mRNAs (Figure 1C right), suggesting adaptation to the microgravity environment. This phenomenon may be due to the constant reduction in the abundance of these mRNAs (Figure 1C middle), which allows long-term suppression of mitochondrial protein production. On the other hand, this adaptation did not occur in the simulated microgravity condition condition at least for 72 h (Figures 2C and S7B). This difference may be due to the lesser magnitude of the effect caused by simulated microgravity than by bona fide microgravity in the ISS (compare the y-axis scales in Figures 1C and 2C), probably owing to the “time-averaged” microgravity in the 3D clinostat.

By small-scale chemical screening, we outlined the signaling cascade involved in cell adhesion to the mitochondrial outer membrane (Figure 6D). Earlier studies indicated that PAK1 phosphorylates BAD in two pathways: directly or indirectly through Raf1 ^48^. Since we could not detect a significant effect of the Raf1 inhibitor (Figure S7C-E), direct phosphorylation by PAK1 may be a major pathway under our conditions, although our data did not exclude a variable balance of the direct and indirect pathways across cells. Although the pan-Bcl-2 family protein inhibitors obatoclax and TW-37 blocked laminin-activated mitochondrial translation (Figure 4F), the same phenotype was not observed for the compounds specific for each Bcl-2 family protein (Figure S7F-H), suggesting that Bcl-2 family proteins perform redundant functions in enhancing mitochondrial translation.

This study provides a framework for mitochondrial translation activation primed by laminin–integrin but simultaneously raises new questions. It remains unclear how the Bcl-2 family proteins on the mitochondrial outer membrane transmit signals to mtFAS in the mitochondrial matrix. The prosurvival effect of Bcl-2 family proteins relies on their binding to BAX to suppress the oligomerization of BAX, a driver of the apoptotic cascade. However, the function of the Bcl-2 family in activating mitochondrial translation may not be based on this conventional mechanism, since neither the BAX activator nor the inhibitor impacted mitochondrial translation (Figure S7I-K).

Furthermore, how malonylation in the translation machinery affects the process of initiation and elongation should be addressed in future studies. Malonylation may affect the interaction between mitoribosomes and other factors. For example, malonylated lysines in uL12m, which consists of the L7/L12 stalk ^87^, were located at the interface with mtIF2, mtEFG1, and mtEFG2 (Figures 5G, S6C, and S6E). Similarly, the interaction between K191 of uL11m and E562 of mtEGF1 ^88^ could be interfered with malonylation of K191 and K192 of uL11m (Figure 5G).

Our study elucidates the unique translational response of bioenergetic powerhouses to external gravitational and mechanical forces.

## Acknowledgments

We thank all the members of the Iwasaki laboratory and Afshin Beheshti (National Aeronautics and Space Administration, NASA) for constructive discussion, technical help, and critical reading of the manuscript. We are also grateful to Yasuhiro Nakamura (Japan Aerospace Exploration Agency, JAXA), Toru Shimazu (Japan Space Forum), Hiromi Sano (Japan Manned Space Systems Corporation, JAMSS), and Ikuko Osada (JAMSS) for administrative and technical help with the spaceflight experiments (termed “Ribosome Profiling”). The ground operation for the spaceflight experiments was supported by the JAXA Flight Control Team and Payload Flight Control Team. The experiments in the ISS were conducted by Soichi Noguchi. FACS analysis was supported by Kenji Ohtawa from the Support Unit for Bio-Material Analysis, RIKEN CBS Research Resources Division. Microscopic analysis was supported by RIKEN CBS-Olympus Collaboration Center. A part of the deep sequencing analysis was conducted by the Vincent J. Coates Genomics Sequencing Laboratory at UC Berkeley, supported by the NIH S10 OD018174 Instrumentation Grant. This work used the supercomputer HOKUSAI SailingShip in RIKEN. Mouse 3T3 cells were gifts from Shinichi Nakagawa. S.I. was supported by the Japan Society for the Promotion of Science (JSPS) (a Grant-in-Aid for Young Scientists [A], JP17H04998; a Challenging Research [Exploratory], JP19K22406; a Grant-in-Aid for Scientific Research [B], JP23H02415), the Ministry of Education, Culture, Sports, Science and Technology (MEXT) (a Grant-in-Aid for Transformative Research Areas [B] “Parametric Translation”, JP20H05784), the Japan Agency for Medical Research and Development (AMED) (AMED-CREST, 23gm1410001), the Gushinkai Foundation, and RIKEN (“Biology of Intracellular Environments”). T.W. was supported by The Graduate School of Frontier Sciences, The University of Tokyo (the Challenging New Area Doctoral Research Grant, C2205) and by JSPS (a Grant-in-Aid for JSPS Fellows, JP23KJ0444). Y.K. was supported by JSPS (a Grant-in-Aid for JSPS Fellows, JP20J10665). K.N. was supported by The Graduate School of Frontier Sciences, The University of Tokyo (the Challenging New Area Doctoral Research Grant, C2205) and JSPS (a Grant-in-Aid for JSPS Fellows, JP22J23099 and JP22KJ1154). Y.S. was supported by JSPS (a Grant-in-Aid for Early-Career Scientists, JP21K15023), MEXT (a Grant-in-Aid for Transformative Research Areas [A] “Multifaceted Proteins”, JP21H05734), AMED (AMED-PRIME, JP23gm6910005h0001), the Exploratory Research Center on Life and Living Systems (ExCELLS, 23EX601), and RIKEN (Pioneering Projects “Biology of Intracellular Environments”). T. Tsuboi acknowledges support from the Jilin Fuyuan Guan Food Group Co., Ltd, the Science, Technology, Innovation Commission of Shenzhen Municipality (WDZC20220811144737001), startup funds from Tsinghua SIGS, and the National Key R&D Program of China (2023YFA0914300). Y.H. was supported by AMED (JP19dm0207082 and JP21wm0525015) and JSPS (a Grant-in-Aid for Transformative Research Areas [A] “New cross-scale biology”, JP 22H05532). We also thank the following fellowships: the RIKEN Junior Research Associate Program for Y.K. and H.S.; the JSPS Research Fellow [DC1] for K.N.; the JSPS Research Fellow [DC2] for Y.K., T.W. and T. Tsubaki; JST SPRING [JPMJSP2108] for T.W. and T. Tsubaki; and the ANRI fellowship for T.W.).

## Author Contributions

Conceptualization, T.W., Y.K., T. Yamamori, T. Yamazaki, A. Higashibata, and S.I.; Methodology, T.W., Y.K., M.M., T. Tsubaki, M.L., K.N., A.H.K., H.S., T. Yamamori, T. Yamazaki, A. Higashibata, T. Tsuboi, Y.H., N.T.-T., T. Saito, A. Higashitani, Y.S., and S.I.; Formal analysis, T.W., Y.K., M.M., T. Tsubaki, M.L., K.N., A.H.K., H.S., and T. Tsuboi; Investigation, T.W., Y.K., M.M., T. Tsubaki, M.L., K.N., and H.S.; Resources, A. Higashibata and A. Higashitani; Writing – Original Draft, T.W., Y.K., and S.I.; Writing – Review & Editing, T.W., Y.K., M.M., T. Tsubaki, M.L., K.N., A.H.K., H.S., T. Yamamori, T. Yamazaki, A. Higashibata, T. Tsuboi, Y.H., N.T.-T., T. Saito, A. Higashitani, Y.S., and S.I.; Visualization, T.W. and S.I.; Supervision, T. Yamamori, T. Yamazaki, A. Higashibata, T. Tsuboi, Y.H., N.T.-T., T. Saito, A. Higashitani, Y.S., and S.I.; Funding Acquisition, T.W., Y.K., K.N., T. Tsuboi, Y.H., Y.S., and S.I.

## Experimental Procedures

### Cell culture

HEK293 cells (ATCC, CRL-1573), C2C12 cells (ATCC, CRL-1772), 3T3 cells (kind gifts from Shinichi Nakagawa’s laboratory), naïve HAP1 cells (Horizon Discovery, C631), *BAD* KO HAP1 cells (Horizon Discovery, 29-bp deletion, HZGHC005959c002), naïve MEF cells (ATCC, CRL-2991), and *OPA1 KO* MEF cells (ATCC, CRL-2995) were cultured in DMEM (high glucose, GlutaMAX Supplement, Thermo Fisher Scientific, 10566016) supplemented with 10% FBS (Sigma‒Aldrich) (for HEK293, C2C12, and 3T3), DMEM supplemented with 10% FBS, 1 × Na-pyruvate (Nakalai, 06977-34) and 0.05 g/L uridine (TCI, U0020) (for MEF), or IMDM (GlutaMAX Supplement, Thermo Fisher Scientific, 31980030) supplemented with 10% FBS (Sigma‒Aldrich) (for HAP1) at 37°C with 5% CO_2_, according to the manufacturer’s instructions. The absence of *Mycoplasma*-free culture was confirmed by an e-Myco VALiD Mycoplasma PCR Detection Kit (iNtRON Biotechnology). A disposable cultivation chamber (DCC) (JAXA and Chiyoda Corporation) ^89^ was used to culture the cells on a 3D clinostat (5.5 rpm on the X axis and 6.5 rpm on the Y axis; AES) and a centrifuge (AES). The DCC was filled with medium to prevent shear stress caused by air bubbles.

### Compounds

Cells were treated with the following compounds at the indicated concentrations: cycloheximide (Sigma‒Aldrich, C4859, 100 μg/ml), chloramphenicol (FUJIFILM Wako Pure Chemical Corporation, 030-19452, 100 μg/ml), anisomycin (Alomone Labs, A-520, 100 μg/ml), RGDS peptide (Tocris, 3498, 10 μg/ml), NSC23766 (Tocris, 2161, 200 μM), ZCL278 (Tocris, 4794, 100 μM), Rhosin (Tocris, 5003, 50 μM), BAY-293 (Tocris, 6857, 1 μM), LY294002 (Sigma‒Aldrich, L9908, 10 μM), SP600125 (Sigma‒Aldrich, S5567, 20 μM), IPA-3 (Tocris, 3622, 5 μM), CK666 (Tocris, 3950, 25 μM), obatoclax (Selleckchem, S1057, 2 μM), TW-37 (Selleckchem, S1121, 10 μM), valinomycin (Tocris, 3373, 2 μM), sorafenib (Selleckchem, S1040, 1 μM), GW5074 (Tocris, 1381, 5 μM), ZM336372 (Tocris, 1321, 10 μM), venetoclax (Selleckchem, S8048, 10 μM), A-1331852 (Selleckchem, S7801, 10 μM), S63845 (Selleckchem, S8383, 5 μM), BAI1 (Tocris, 2160, 1 μM), BTSA1 (Selleckchem, S8650, 20 μM), BAM7 (Tocris, 4810, 20 μM), C75 (Tocris, 4810, 50 μM), nicotinamide (Sigma‒Aldrich, N3376, 10 mM), NRD167 (Selleckchem, S9903, 10 μM), and 3-TYP (Selleckchem, S8628, 10 μM). Chloramphenicol was dissolved in 70% ethanol, nicotinamide was dissolved directly in the medium, and the other compounds were dissolved in dimethyl sulfoxide (DMSO).

### Spaceflight experiments

DCCs were pretreated with laminin overnight as described below. Then, 5 × 10^5^ HEK293 cells were seeded in each DCC and incubated overnight. The medium was exchanged for CELLBANKER 1 plus (TaKaRa), and the cells were stored at −80°C. The frozen cells were then sent to the ISS by Cygnus NG-14 in October 2020. In the Japanese Experiment Module KIBO in the ISS, the samples were thawed, exchanged for DMEM, high glucose, GlutaMAX Supplement (Thermo Fisher Scientific) supplemented with 10% FBS using a Pre-Fixation Kit-III (PFK-III, JAXA) ^89^ and cultured under centrifugation at 1 × *g* in the Cell Biology Experiment Facility (CBEF) ^90^ at 37°C with 5% CO_2_ and 80±5% RH for 24 h. Subsequently, the cells were cultured under microgravity for 24 and 48 h, while the control cells were maintained at 1 × *g*. For chemical fixation, the cell culture media were replaced with CELLBANKER 1 plus containing 100 µg/ml cycloheximide and 100 µg/ml chloramphenicol by PFK-III. The samples were subsequently stored in a Minus Eighty degree Celsius Laboratory Freezer for ISS (MELFI) at −95°C. The samples were returned to the ground by SpaceX CRS-22 (SpX-22) in July 2021 and shipped to the laboratory for ribosome profiling and RNA-seq.

Space-flown *C. elegans* samples prepared for earlier experiments ^16^ were used in this study.

### Animals

All animal experiments were authorized and approved by the Animal Care and Use Committee of The University of Tokyo. Adult male (7 weeks old) C57BL/6J mice were purchased from Sankyo Lab Service Corporation, housed in the facility under the supervision of the IACUC at 18 to 22°C with a 12-h:12-h light–dark cycle, and allowed free access to food and water.

#### Minimized mechanical stress (MMS)

The MMS model was developed by one of the coauthors and will be published elsewhere. First, we applied adhesive tape to the tails of 8-week-old mice. The knee joints of the mice were then kept immobilized using plastic cylinders. The feet of the mice were also taped to prevent the plastic cylinders from sliding off. Additional adhesive tape was applied between the rear limbs so that the mice could not bite the adhesive tape around their tail. Following joint fixation, the tails of the mice were suspended by attaching a spring clip to the adhesive tape. The spring clip was connected to a piano wire that was attached to the cage lid so that the mice could move in their cages. Food was left on the floor, and a water bottle was fixed to the wall with double-sided tape for easy access.

Mice were sacrificed at 1, 3, and 14 d after MMS started. For the control, age-matched mice were harvested (8 weeks old and 10 weeks old). The soleus muscles were dissected, and the wet weights were measured. The muscles were immediately flash-frozen in liquid nitrogen and stored at −80°C.

### Ribosome profiling and RNA-Seq

#### Lysate preparation

##### Spaceflight samples, HEK293

The cells in a DCC were thawed, washed with ice-cold PBS, lysed with 500 µl of lysis buffer (20 mM Tris-HCl pH 7.5, 150 mM NaCl, 5 mM MgCl_2_, 1 mM dithiothreitol [DTT], 1% Triton X-100, 100 µg/ml cycloheximide, and 100 µg/ml chloramphenicol), and triturated ten times in a syringe with a 30-gauge needle (NIPRO Corporation). The lysate was incubated with 25 U/ml TURBO DNase (Thermo Fisher Scientific) and clarified by centrifugation at 20,000 × *g* for 10 min at 4°C.

##### Spaceflight samples, C. elegans

Frozen nematode cultures (1.5-4.5 ml) were thawed in 10 ml of wash buffer (20 mM Tris-HCl pH 7.5, 150 mM NaCl, 5 mM MgCl_2_, 100 µg/ml cycloheximide, and 100 µg/ml chloramphenicol). The adult worms were filtered through a 30-µm ÜberStrainer (pluriSelect Life Science) and washed three times with wash buffer. Then, the nematodes were pelleted by centrifugation at 1,600 × *g* for 1 min at 4°C, resuspended in 600 µl of lysis buffer, and dripped into liquid nitrogen. After pulverization using a Multi-beads Shocker (Yasui Kikai) at 2800 rpm for 10 sec, the lysate was thawed at 4°C, incubated with 25 U/ml Turbo DNase (Thermo Fisher Scientific), and clarified by centrifugation at 20,000 × *g* for 10 min at 4°C.

##### Simulated microgravity samples

Cells in a DCC were washed with PBS and lysed with 500 µl of lysis buffer. After 25 U/ml TURBO DNase treatment (Thermo Fisher Scientific), the lysate was clarified by centrifugation at 20,000 × *g* for 10 min at 4°C.

##### Mouse soleus muscles

Frozen soleus muscles were pulverized with 600 µl of frozen lysis buffer containing cOmplete, EDTA-free Protease Inhibitor Cocktail (Roche) and 1000 U/ml RNase inhibitor, Murine (New England Biolabs [NEB]), using a Multi-beads Shocker (Yasui Kikai) at 3000 rpm for 30 sec for 2 cycles. After thawing at 4°C, the lysate was incubated with 25 U/ml Turbo DNase (Thermo Fisher Scientific) and clarified by centrifugation at 20,000 × *g* for 10 min at 4°C.

#### Library preparation for ribosome profiling (standard Ribo-Seq and Thor-Ribo-Seq)

For the simulated microgravity samples, ribosome profiling library preparation was performed as previously described ^91^. The other libraries were generated by the Thor-Ribo-Seq protocol as described earlier ^25^.

The lysate was treated with 20 U of RNase I (LGC Biosearch Technologies) at 25°C for 45 min. A sucrose cushion was used to collect ribosomes. The RNA was run on a 15% UREA PAGE gel. RNA fragments ranging from 17 to 34 nucleotides (nt) were excised from the gel, dephosphorylated, and ligated with linkers. rRNAs were depleted with the Ribo-Zero Gold rRNA Removal Kit (Human/Mouse/Rat) (Illumina) for human and mouse samples and with *Caenorhabditis elegans* Ribo-Seq riboPOOLs (siTOOLs Biotech) for nematode samples. For the simulated microgravity samples, cDNA was reverse-transcribed, circular-ligated, and PCR-amplified. For the other samples, after hybridization of an oligonucleotide to the T7 promoter region of the linker, complementary RNAs were transcribed by a T7-Scribe Standard RNA IVT Kit (CELLSCRIPT) and then ligated to the second linker. cDNA was reverse-transcribed and PCR-amplified. The DNA libraries were sequenced on a HiSeq 4000 platform in 50-bp single-read mode (simulated microgravity experiments) or on a HiSeq X Ten platform in 150-bp paired-end mode (the other experiments).

#### Library preparation for RNA-Seq

For RNA-seq, total RNA was extracted from the same lysate used for ribosome profiling with TRIzol LS (Thermo Fisher Scientific) and a Direct-zol RNA Microprep Kit (Zymo Research). rRNA depletion was conducted with the Ribo-Zero Gold rRNA Removal Kit (Human/Mouse/Rat) (Illumina) for human and mouse samples and with *Caenorhabditis elegans* Ribo-Seq riboPOOLs (siTOOLs Biotech) for nematode samples.

For HEK293 cells in spaceflight experiments and mouse soleus muscles, libraries were prepared with a SMARTer Stranded Total RNA-Seq Kit v3 - Pico Input (TaKaRa). The other RNA-Seq libraries were generated with a TruSeq Stranded mRNA Library Prep Kit (Illumina). The DNA libraries were sequenced on a HiSeq 4000 platform in 50-bp single-read mode (s-µg experiments) or on a HiSeq X Ten platform in 150-bp paired-end mode (the other experiments).

### MitoIP-Thor-Ribo-Seq

#### Lysate preparation

Lysate was prepared as previously reported ^77^. Cells in 15-cm dishes were washed with ice-cold PBS and lysed with 1200 µl of hypotonic buffer (10 mM HEPES-KOH pH 7.5, 10 mM KCl, 1.5 mM MgCl_2_, 1 mM DTT, 100 µg/ml cycloheximide, and 100 µg/ml chloramphenicol). The mitochondrial fraction in the lysate was purified through immunoprecipitation with an anti-TOM22 antibody from the Mitochondria Isolation Kit, Human (Miltenyi Biotec), according to the manufacturer’s instructions with minor modifications. Then, the eluted mitochondria were pelleted by centrifugation at 7,000 × *g* for 10 min at 4°C and resuspended in modified lysis buffer (20 mM Tris-HCl pH 7.5, 150 mM NaCl, 15 mM MgCl_2_, 1 mM DTT, 1% Triton X-100, 100 µg/ml cycloheximide, 100 µg/ml chloramphenicol, and 2.5 U/ml TURBO DNase). The lysate was clarified by centrifugation at 20,000 × *g* for 10 min at 4°C for library preparation.

#### Library preparation

Library preparation was performed as previously described ^77^. The lysate was incubated with 40 U of RNase I (LGC Biosearch Technologies) in a 50-µl reaction (scaled up by the modified lysis buffer) at 25°C for 45 min. After sucrose cushion ultracentrifugation and Ribo-FilterOut ^92^, RNA fragments ranging from 17 to 50 nt were collected.

From linker-ligated RNA fragments, rRNAs were depleted with Human Ribo-Seq riboPOOL (siTOOLs Biotech). Then, an oligonucleotide was hybridized to the T7 promoter region of the linker to generate the dsDNA T7 promoter. Complementary RNAs were transcribed with a T7-Scribe Standard RNA IVT Kit (CELLSCRIPT). After ligation of the second linker, cDNA was reverse-transcribed and PCR-amplified. The DNA libraries were sequenced on a HiSeq X Ten in 150-bp paired-end mode.

## Data analysis

The deep sequencing data were processed as described previously ^27,93^, with modifications. For ribosome profiling with paired-end sequencing, base correction was performed by Fastp (version 0.21.0) ^94^, and read 1 was used for downstream analysis. After read quality filtering and adapter trimming by Fastp, all reads aligned to noncoding RNAs (rRNAs, tRNAs, mt-rRNAs, mt-tRNAs, snRNAs, snoRNAs, and miRNAs) were removed using STAR (version 2.7.0a) ^95^. The reads were subsequently mapped to the corresponding nuclear genomes (human, hg38; mouse, mm10; *C. elegans*, WBcel235) and custom-made mitochondrial transcript sequences by STAR. UMI suppression in Thor-Ribo-Seq was performed using UMItools (version 1.1.2) ^96^. For ribosome profiling, ribosomal A-site position offsets from the 5′ end of reads were empirically estimated along the read length, as summarized in Table S2. For RNA-Seq, the offset was set to 15. To count reads from CDSs, reads assigned to the first and last 5 amino acids were excluded. The relative enrichment of reads across CDSs in ribosome profiling and RNA-Seq were calculated by the DESeq2 package ^97^. Additionally, translation efficiency was calculated by a generalized linear model with the same package. Gene Ontology analysis was performed with DAVID (https://david.ncifcrf.gov/home.jsp) ^98,99^. Mitocarta3 genes were defined at https://www.broadinstitute.org/files/shared/metabolism/mitocarta/human.mitocarta3.0.html) ^100^. The mtUPR target genes were defined in an earlier study ^36^.

The global mitochondrial elongation rate was calculated as previously reported ^75^. Smoothed metagene profiles were generated from the mitochondrial transcripts, excluding *MT-ATP6* and *MT-ND4* due to the bicistronic nature. The mean read density from 100 to 200 codons was normalized to 1.

### Laminin precoating

DCCs, 6-well plates, 12-well plates, and 96-well plates were precoated with 0.25 µg/cm^2^ (or dilutions as indicated in the figures) laminin (Easy iMatrix-511 silk, MATRIXOME) overnight at 4°C before cell seeding.

### Cell adhesion assay

The wells of a 96-well plate were precoated with laminin as described above. One hundred microliters of HEK293 cell suspension (5 × 10^5^ cells/ml) was seeded and incubated for 1 h in a CO_2_ incubator. After washing out the unattached cells with PBS, the dish-attached cells were fixed with 4% (w/v) paraformaldehyde in PBS for 20 min, permeabilized with 0.1% Triton X-100 in PBS, and stained with 0.2 μM CellTag 700 (LI-COR Biosciences) for 1 h. Images were acquired by an Odyssey CLx (LI-COR Biosciences) with an IR 700-nm channel.

### On-gel mito-FUNCAT

On-gel mito-FUNCAT was conducted as described previously ^31^. Unless otherwise noted, cells were seeded on dishes precoated with or without laminin and incubated for 12 h with the indicated compounds before cell harvesting. Nascent peptides were labeled with 50 μM HPG (Jena Bioscience) in methionine-free medium (DMEM, high glucose, no glutamine, no methionine, no cystine [Thermo Fisher Scientific], supplemented with 48 µg/ml L-cysteine and 4.08 mM L-alanyl-L-glutamine) containing 100 mg/ml anisomycin for 3 or 4 h. Cells were washed with ice-cold PBS and lysed with FUNCAT lysis buffer (20 mM Tris-HCl pH 7.5, 150 mM NaCl, 5 mM MgCl_2_, and 1% Triton X-100). After clarification by centrifugation at 20,000 × *g* for 10 min at 4°C, the supernatants were labeled with 50 μM IRdye800CW Azide (LI-COR Biosciences) via a Click-it Cell Reaction Buffer Kit (Thermo Fisher Scientific), according to the manufacturer’s instructions. The proteins were separated on SDS–PAGE gels. Images were acquired with an Odyssey CLx (LI-COR Biosciences) with an IR 800-nm channel. For signal normalization, the gels were stained with EzStain AQua (ATTO) or GelCode Blue Safe Protein Stain (Thermo Fisher Scientific) to detect total proteins, and images were acquired with an Odyssey CLx with an IR 700-nm channel. The signals on the gels were quantified with Image Studio (LI-COR Biosciences, version 5.2).

For the measurement of total protein synthesis, anisomycin was omitted from the experiments.

For knockdown experiments, cells were transfected with 5 μM siRNA targeting FAK (Horizon Discovery, L-003164-00-0005) or control siRNA (Horizon Discovery, D-001206-13-20) using TransIT-X2 Reagent (Mirus Bio, MIR6003) in 10-cm dishes and incubated for 24 h. Then, the cells were reseeded on 12-well dishes with or without laminin coating as described above, cultured for an additional 24 h, and used for on-gel mito-FUNCAT experiments.

### Mito-FUNCAT FACS

Mito-FUNCAT FACS was performed as described previously ^31^. HAP1 cells were cultured in methionine-free medium supplemented with 100 µM HPG and 100 mg/ml anisomycin for 3 h before cell harvesting. After washing with PBS, the cells were dissociated from the dish with 0.05% trypsin. The collected cells were washed, prepermeabilized with 0.0005% digitonin in mitochondrial protective buffer (10 mM HEPES-KOH pH 7.5, 300 mM sucrose, 10 mM NaCl, and 5 mM MgCl_2_) for 5 min at room temperature, fixed in 4% paraformaldehyde (PFA) for 15 min at 4°C, and permeabilized with 0.1% (v/v) Triton X-100 in PBS for 5 min at room temperature. Subsequently, nascent peptides within mitochondria were labeled with 1 μM azide-conjugated Cy3 (Jena Bioscience) with a Click-it Cell Reaction Buffer Kit (Thermo Fisher Scientific). Mitochondria were immunostained with Alexa Fluor 647 Anti-TOMM20 antibody (Abcam, ab209606, 1:100) in Intercept Blocking Buffer (LI-COR Biosciences) for 1 h at 4°C. The cells were washed three times with PBS and then analyzed via flow cytometry (BD FACSAria II, BD Biosciences).

### Mitochondrial DNA content measurement

Total DNA was extracted from 1 × 10^6^ cells using NucleoSpin DNA RapidLyse (MACHEREY-NAGEL). qPCR was performed with primers targeting the ND2 locus in the mitochondrial genome (5′-TGTTGGTTATACCCTTCCCGTACTA-3′ and 5′-CCTGCAAAGATGGTAGAGTAGATGA-3′) and those targeting the Alu repeat sequence in the nuclear genome (5′-CTTGCAGTGAGCCGAGATT-3′ and 5′-GAGACGGAGTCTCGCTCTGTC-3′) ^101^ with TB Green Premix Ex Taq II (TaKaRa) on a Thermal Cycler Dice Real Time System II (TaKaRa). The relative mitochondrial DNA content was normalized to the nuclear-genome content by the ΔΔCt method.

### Microscopic analysis

The cells were cultured in a glass-bottomed DCC (AES), washed with PBS, fixed with 4% (w/v) paraformaldehyde in PBS for 15 min at room temperature, and subsequently incubated with methanol for 10 min at −20°C. After washing twice with PBS, the cells were incubated with Intercept Blocking Buffer (LI-COR Biosciences) containing 0.2% Triton X-100 for 60 min at room temperature. Then, the mitochondria were immunostained in Intercept Blocking Buffer with 0.2% Triton X-100 and Alexa Fluor 647 Anti-TOMM20 antibody (Abcam, ab209606, 1:100) for 1 h at room temperature. The cells were washed with TBS containing 0.2% Triton X-100 three times and then incubated with VECTASHIELD Mounting Medium (VECTOR LABORATORIES, INC., H-1500) overnight at 4°C in the dark. Images were obtained using an FV3000 confocal microscope (Olympus) with a 60× objective lens (Olympus Japan, UPLXAPO60XO) via the processes described below.

### Morphological quantification of mitochondria

Image analysis was performed using MitoGraph (version 3.0), a fully automated C++ program for 3D mitochondrial structure analysis (https://github.com/vianamp/MitoGraph) ^102,103^. We first isolated 3D images of individual cells from the raw images using the FIJI (https://fiji.sc) macro to extract cellular trends and then applied them as inputs to the MitoGraph software. For the MitoGraph parameter, we used 0.414 for the flag-xy and 0.3 for the flag-z.

### Mitochondrial membrane potential assessment

The mitochondrial membrane potential was measured by flow cytometry as described previously ^104^. The cells were incubated with 200 nM MitoTracker Red CMXRos (Thermo Fisher Scientific) and 150 nM MitoTracker Green FM (Thermo Fisher Scientific) at 37°C for 25 min and then washed with PBS supplemented with 5% FBS. The cells were dissociated from the dish with 0.05% trypsin, neutralized, collected by centrifugation at 300 × *g* for 3 min, resuspended in PBS supplemented with 5% FBS, and then subjected to flow cytometry (BD FACSAria II, BD Biosciences).

### Western blotting

Anti-β-actin (MEDICAL & BIOLOGICAL LABORATORIES [MBL], M177-3, 1:1000), anti-GAPDH (Cell Signaling Technology [CST], 2118, 1:1000), anti-FAK (CST, 3285, 1:1000; Thermo Fisher Scientific, AHO0502, 1:1000), anti-p-FAK^Y397^ (CST, 3283, 1:200), anti-TOMM20 (CST, 42406, 1:1000), anti-RAC1 (CST, 8631, 1:200), anti-PAK1 (CST, 2602, 1:1000), anti-p-PAK1^S144^/p-PAK2^S141^ (CST, 2606, 1:200), anti-BAD (CST, 9292, 1:200), anti-lipoic acid (Abcam, ab58724, 1:1000), anti-DLAT (Abcam, ab172617, 1:1000), anti-MRPS22/mS22 (Thermo Fisher Scientific, PA5-52249, 1:1000), and anti-Mal-K (PTM Bio,PTM-201, 1:1000) were used as primary antibodies. IRDye800CW anti-rabbit IgG (LI-COR Biosciences, 926-32211, 1:10000), IRDye680RD anti-rabbit IgG (LI-COR Biosciences, 925-68071, 1:10000), and IRDye680RD anti-mouse IgG (LI-COR Biosciences, 925-68070, 1:10000) were used as secondary antibodies. Images were obtained with an Odyssey CLx (LI-COR Biosciences) and quantified with Image Studio (LI-COR Biosciences, version 5.2).

### Isolation of RAC1 in the GTP-bound form

RAC1 in the GTP-bound form was isolated with an Active Rac1 Detection Kit (CST, 8815) according to the manufacturer’s instructions. Anti-RAC1 Western blotting was performed as described above.

### Mitochondrial purification

Mitochondria were isolated with a Mitochondria Isolation Kit, Human (Miltenyi Biotec), according to the manufacturer’s instructions with minor modifications. The cells were washed with ice-cold PBS and lysed with hypotonic buffer (10 mM HEPES-KOH pH 7.5, 10 mM KCl, 1.5 mM MgCl_2_, and 1 mM DTT). The cell lysate was incubated with TOM22 beads for 60 min at 4°C. After washing, the mitochondrial fraction was eluted from the beads, pelleted by centrifugation at 7,000 × *g* for 10 min at 4°C, resuspended in modified lysis buffer (20 mM Tris-HCl, pH 7.5, 150 mM NaCl, 15 mM MgCl_2_, 1 mM DTT, and 1% Triton X-100), and then centrifuged at 20,000 × *g* for 10 min at 4°C. The supernatant was subjected to Western blotting.

### Cell viability assay

Cell viability was monitored as described previously ^105^. Cells were seeded on 96-well plates, treated with the indicated compounds for 12 h, and then incubated with RealTime-Glo MT Cell Viability Assay reagent (Promega) for 30 min. The luminescence was quantified by the GloMax Navigator System (Promega).

### Oxygen consumption rate (OCR)

After the cells were cultured on Seahorse XFe96 cell culture microplates (Agilent), the medium was replaced with Seahorse XF DMEM (Agilent) containing 1 mM pyruvate, 2 mM glutamine, and 10 mM glucose and then incubated at 37°C for 1 h. The OCR was monitored on a Seahorse XFe96 Analyzer (Agilent) throughout sequential injections of 0.5 μM oligomycin, 0.5 μM FCCP, and 0.5 μM rotenone/antimycin A into the cell medium, according to the manufacturer’s instructions.

### Metabolomic analysis

The cells on DCCs were cultured under simulated microgravity or standard gravity (at 1 × *g*) for 24 h, washed with 5% (w/w) mannitol (Fujifilm Wako Pure Chemical Corporation), and incubated with methanol (Fujifilm Wako Pure Chemical Corporation) for 30 s. Subsequently, the cells were incubated with 1 mM internal standard solution (Human Metabolome Technologies [HMT]) and collected in 1.5-ml tubes. The samples were centrifuged at 2,300 × *g* for 5 min at 4°C. The proteins were removed from the supernatant by ultrafiltration filters (HMT). Ionic metabolites were comprehensively measured at HMT using capillary electrophoresis time-of-flight mass spectrometry (CE-TOFMS) and capillary electrophoresis triple-quadrupole mass spectrometry (CE-QqQMS).

### Electron microscopy

The cells were fixed with 2.5% glutaraldehyde (Electron Microscopy Sciences) in DMEM for 1 h at room temperature. After washing with 0.1 M phosphate buffer (0.02 M sodium dihydrogenphosphate dihydrate and 0.08 M disodium hydrogenphosphate), the cells were scraped and collected with 0.2% BSA/0.1 M phosphate buffer followed by centrifugation at 820 × *g*. After being embedded in low melting agarose (2% in 0.1 M phosphate buffer, MP Biomedicals), the cell pellets were sectioned at 200-µm thickness with a Leica VT1000S vibratome. The sections were postfixed with 1% OsO_4_ (Electron Microscopy Sciences) and 1.5% potassium ferrocyanide (FUJIFILM Wako Pure Chemical Corporation) in 0.05 M phosphate buffer for 30 min. After being rinsed 3 times with H_2_O, the cells were stained with 1% thiocarbohydrazide (Sigma‒Aldrich) for 5 min. After being rinsed with H_2_O 3 times, the cells were stained with 1% OsO_4_ in H_2_O for 30 min. After being rinsed with H_2_O 2 times at room temperature and 3 times with H_2_O at 50°C, the cells were treated with Walton’s lead aspartate (0.635% lead nitrate [Sigma‒ Aldrich] and 0.4% aspartic acid [pH 5.2, Sigma‒Aldrich]) at 50°C for 20 min. After incubating with an ascending ethanol series (10 min each in 50% on ice, 70% on ice, and 10 min each in 90%, 95% ethanol/H_2_O at room temperature), the sections were rinsed for 10 min with 99.5% ethanol 4 times. Then, the sections were infiltrated with a 1:1 mixture of EPON812 (TAAB) and ethanol for 22 h. After incubating with 100% EPON812 resin for 4 h, the resin was cured at 65°C for 6 d. EPON812 resin was made by mixing 7.5 g of MNA (TAAB), 13.7 g of Epok812 (TAAB), 3.8 g of DDSA (TAAB), and 0.2 g of DMP-30 (TAAB). Resin blocks were trimmed with a TrimTool diamond knife (Trim 45, DiATOME). Fifty-micron-thick ultrathin sections made with a diamond knife (Ultra 45, DiATOME) were collected on a cleaned silicon wafer strip using a Leica Ultramicrotome (UC7). The ultrathin sections were imaged with a scanning electron microscope (JSM-IT800SHL, JEOL). Imaging was done at 1 kV accelerating voltage, 5 kV specimen voltage, 1,280 × 960 frame size, 6-mm working distance, 6.40 × 4.80-μm field of view and 14.1-μs dwell time, using a Scintillator Backscattered Electron Detector in Beam Deceleration mode. The final pixel size was a 5 nm square.

### Reconstituted mammalian mitochondrial translation system

*In vitro* translation of mammalian mitochondria was conducted as reported previously ^75,76^. Unless otherwise specified, the standard translation mixtures (5 µl) contained 50 mM HEPES-KOH pH 7.5, 100 mM potassium glutamate, 11 mM Mg(OAc)_2_, 0.1 mM spermine, 1 mM DTT, 0.15 mM each amino acid (except for methionine and cysteine), 0.05 mM methionine, 0.1 mM cysteine, 1 mM ATP, 1 mM GTP, 20 mM creatine phosphate, 0.01 µg/µL 10-formyl-5,6,7,8-tetrahydrofolic acid, 100 nM creatine kinase, 20 nM myokinase, 15 nM nucleoside-diphosphate kinase, 15 nM pyrophosphatase, 0.5 µM IF-2mt, 1.0 µM IF-3mt, 5 µM EF-Tumt, 1 µM EF-Tsmt, 0.5 µM EF-G1mt, 0.5 µM EF-G2mt, 0.5 µM RF-1Lmt, 0.5 µM RRFmt, 5 μM methionyl-tRNA transformylase (*E. coli* MTF), 0.2 µM 55S ribosome, 0.0225 A_260_ units of yeast aminoacyl-tRNA mixture, and 0.1 µM mRNA. CoA, acetyl-CoA, and malonyl-CoA were dissolved in 20 mM HEPES-KOH pH 7.5 and 100 mM KOAc pH 7.5 and added to the reactions. The reporter mRNA 3×FLAG-Pgk-nLuc ^75,76^ was used. The reaction mixture was incubated at 37°C for 3 h.

The enzymatic activity of the products was assessed with a Nano-Glo Luciferase Assay System (Promega). The 2-µl reaction was stopped by adding 18 µl of stop solution (20 mM HEPES-KOH pH 7.5, 100 mM KOAc pH 7.5, 2 mM Mg(OAc)_2_, and 0.1 mg/ml RNase A) and then mixed with 20 µl of the substrate in a white 96-well half-area plate. After incubating for 17 min at room temperature, the luminescence was detected with a GloMax Multi Detection System (Promega).

### Accession numbers

The standard Ribo-seq, RNA-Seq, and MitoIP-Thor-Ribo-seq data (GSE222998) obtained in this study were deposited in the National Center for Biotechnology Information (NCBI) Gene Expression Omnibus (GEO) database. This study also used the reported data for the retapamulin-assisted mitoribosome run-off assay (GSE237154) ^77^.

---

**Figure S1.**
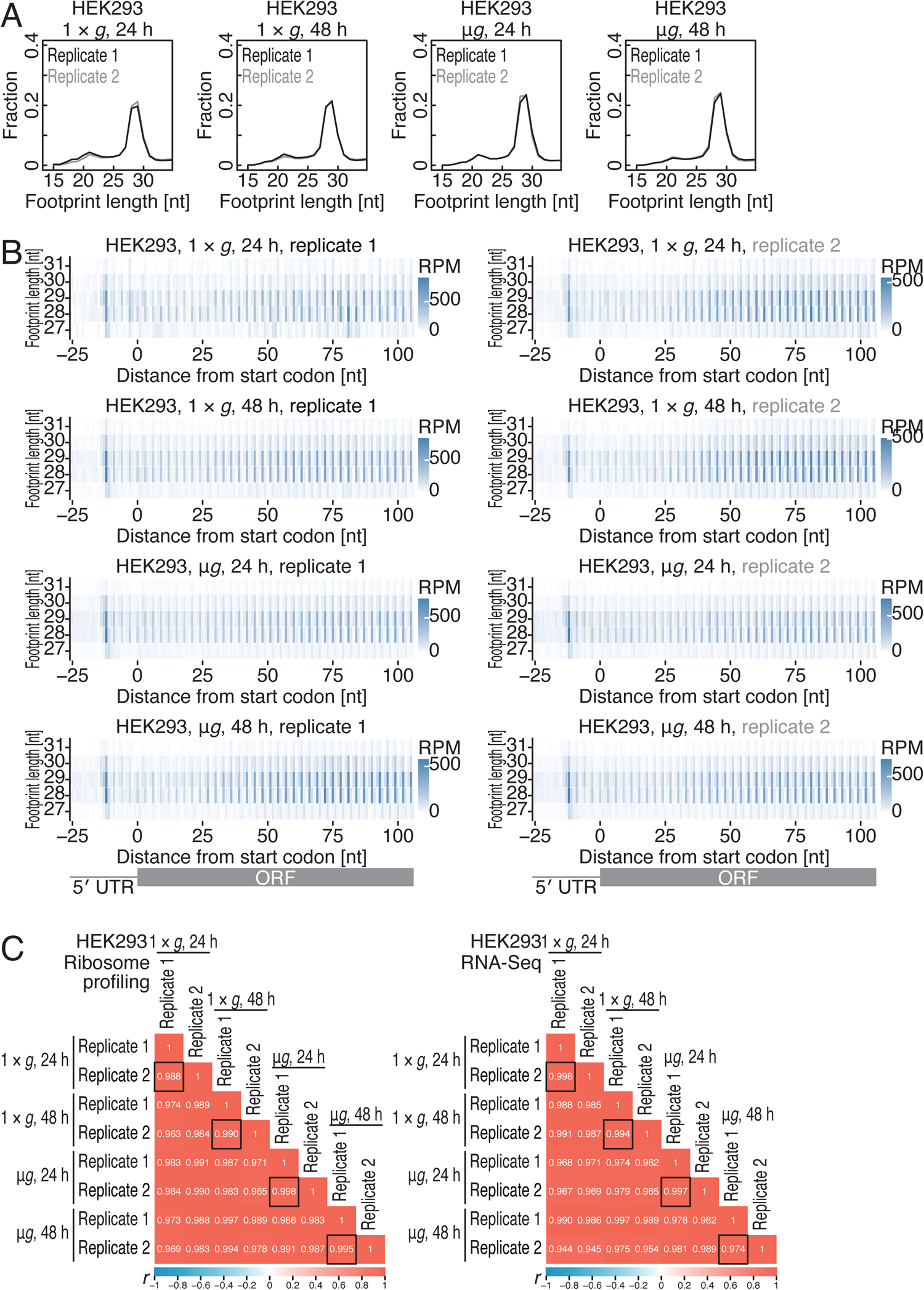
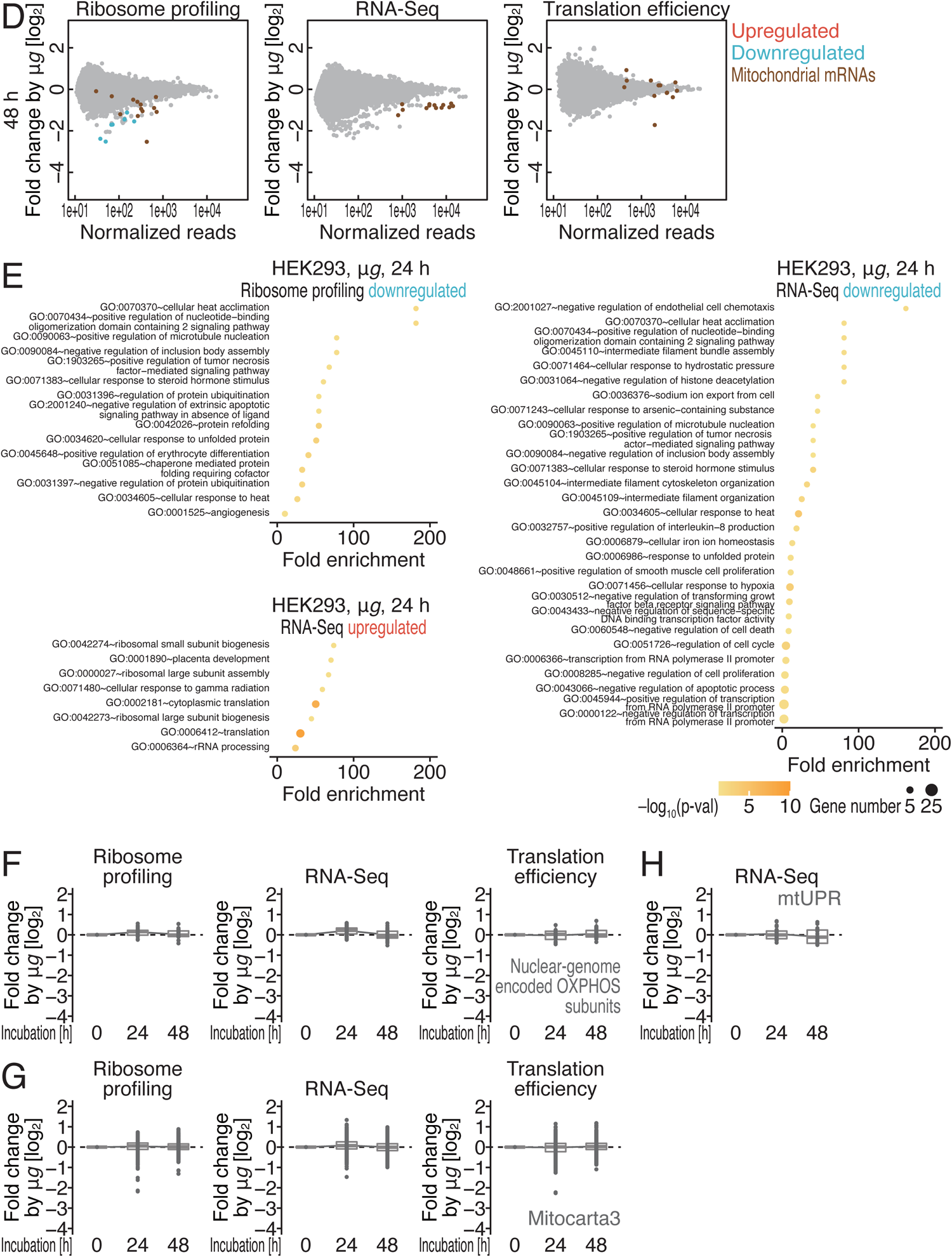
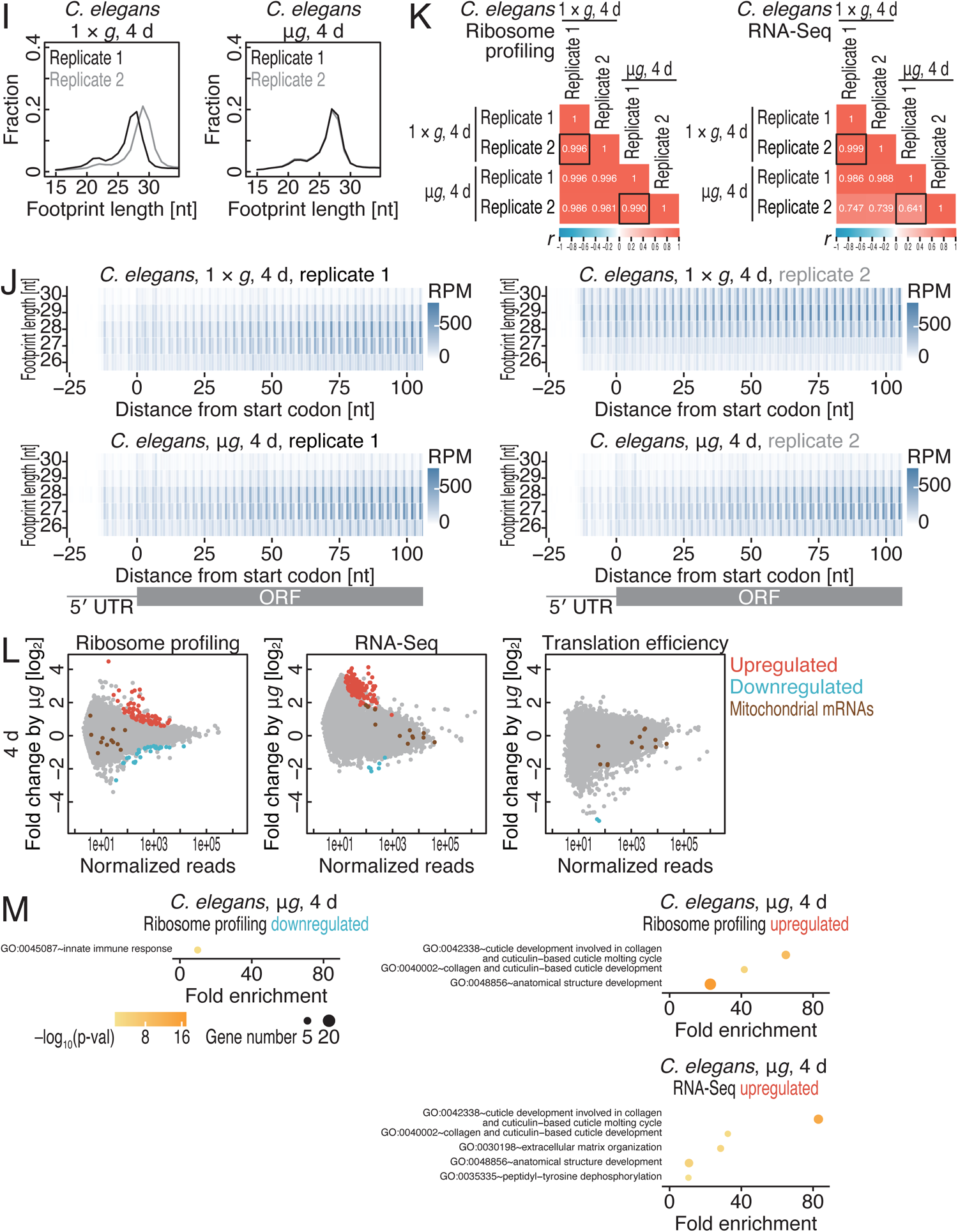
Characterization of ribosome profiling data in spaceflight samples. Related to Figure 1. (A and I) The lengths of ribosome footprints in the indicated samples. (B and J) Metagene plots of ribosome footprints relative to the start codon across the footprint length for the indicated samples. The 5′ end positions of ribosome footprints are depicted. The color indicates the read abundance. RPM, reads per million reads. (C and K) Pearson’s correlation coefficient (*r*, represented in the color as well) among the indicated samples in ribosome profiling and RNA-Seq. The black squares indicate the correlations of the biological replicates. (D and L) MA (M, log ratio; A, mean average) plots for ribosome footprint change (left), RNA abundance change (middle), and translation efficiency change (right) in HEK293 cells after 48-h microgravity (μ*g*) culture (D) and in *C. elegans* after 4-d microgravity culture (L). Significantly altered transcripts (false discovery rate [FDR] < 0.05) and mitochondrial genome-encoded mRNAs are highlighted. (E and M) Gene Ontology (GO) analysis by DAVID ^98,99^ for genes downregulated or upregulated by microgravity in the indicated samples. The color represents statistical significance, and the circle size indicates the number of genes found in the GO category. The p values were calculated by a modified Fisher’s exact test (EASE score). (F and G) Box plots of ribosome footprint change (left), RNA abundance change (middle), and translation efficiency change (right) in nuclear genome-encoded OXPHOS subunits (F) and mitochondria-localized proteins (G) (defined by Mitocarta3 ^100^) over the course of microgravity culture in HEK293 cells. (H) Box plots of the RNA abundance change (middle) of mtUPR target genes (defined by ^36^) over the course of microgravity culture in HEK293 cells. In box plots (F, G, and H), the median (centerline), upper/lower quartiles (box limits), and 1.5× interquartile range (whiskers) are shown.

**Figure S2.**
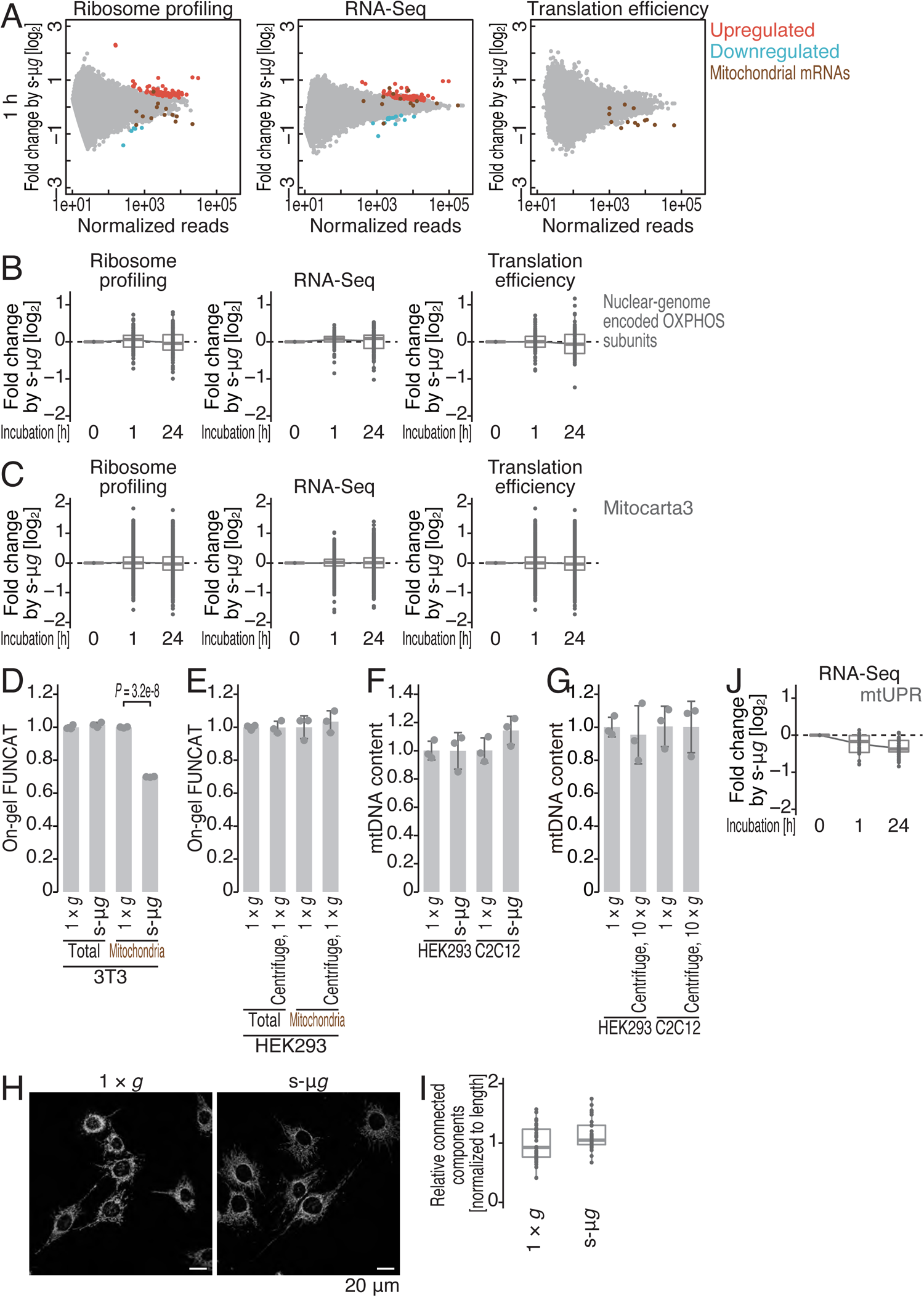
Characterization of ribosome profiling data in HEK293 cells in simulated microgravity. Related to Figure 2. (A) MA plots of ribosome footprint change (left), RNA abundance change (middle), and translation efficiency change (right) after 1-h simulated microgravity culture in HEK293 cells. Significantly altered transcripts (false discovery rate [FDR] < 0.05) and mitochondrial genome-encoded mRNAs are highlighted. (B and C) Box plots of ribosome footprint change (left), RNA abundance change (middle), and translation efficiency change (right) in nuclear-genome-encoded OXPHOS subunits (B) and mitochondria-localized proteins (C) (defined by Mitocarta3 ^100^) over the course of simulated microgravity culture in HEK293 cells. (D and E) On-gel FUNCAT experiments to monitor total and mitochondrial translation in the indicated cell lines and conditions. The cells were cultured under s-μ*g* or 1 × *g* for 24 h before cell harvesting. (F and G) Relative mitochondrial DNA (mtDNA) content in the indicated cell lines and conditions. The cells were cultured under s-μ*g* or 10 × *g* for 24 h before cell harvesting. (H and I) Mitochondria in C2C12 cells were immunostained with an AF647-labeled anti-TOMM20 antibody under the indicated culture conditions. Representative images of immunostaining (H) and quantification of the mitochondrial connected components (I) are shown. Scale bar, 20 μm. (J) Box plots of the RNA abundance change (middle) of mtUPR target genes (defined by ^36^) over the course of simulated microgravity culture in HEK293 cells. For D-G, the data from three replicates (points), the mean values (bars), and the s.d.s (errors) are shown. The p values were calculated by Student’s t test (two-tailed) (D). In box plots (B, C, J, and I), the median (centerline), upper/lower quartiles (box limits), and 1.5× interquartile range (whiskers) are shown.

**Figure S3.**
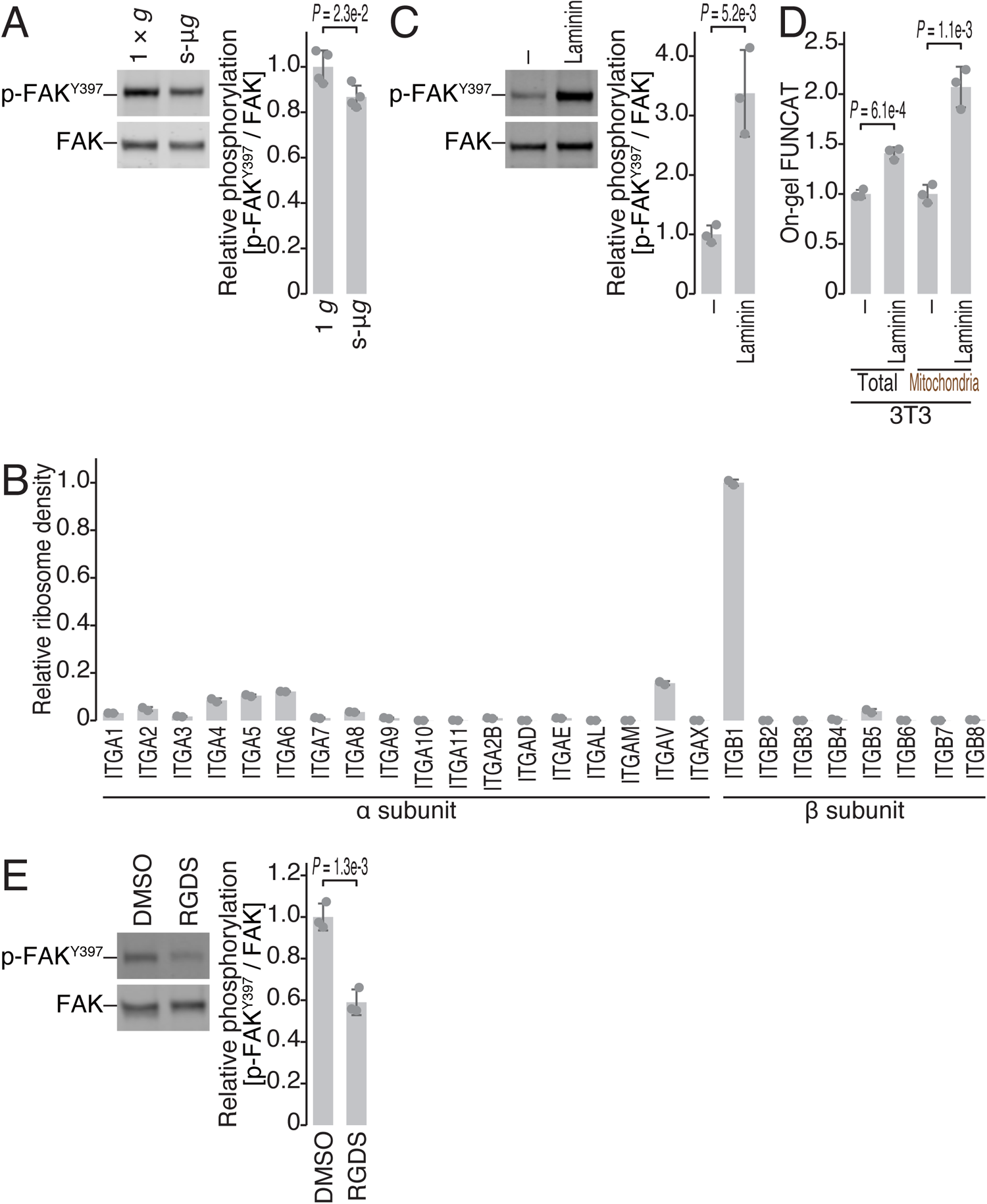
Characterization of the phosphorylation status of FAK. Related to Figure 3. (A, C, and E) Western blotting of total FAK protein and the phosphorylated form (at Y397). The quantified relative amount of phosphorylated FAK is shown. (B) Relative ribosome footprint density (normalized to the average of ITGB1) of the indicated integrin subunit genes. Ribosome profiling data from HEK293 cells cultured at 1 × *g* (Figure 2A-C) were used. (D) On-gel FUNCAT experiments to monitor total and mitochondrial translation in the indicated cell lines and conditions. For A and C-E, the data from replicates (points, n = 4 for A; n = 3 for C-E), the mean values (bars), and the s.d.s (errors) are shown. The p values were calculated by Student’s t test (two-tailed) (A and C-E). For B, the data from two replicates (points) and the mean values (bars) are shown. The experiments were conducted in HEK293 cells unless otherwise noted.

**Figure S4.**
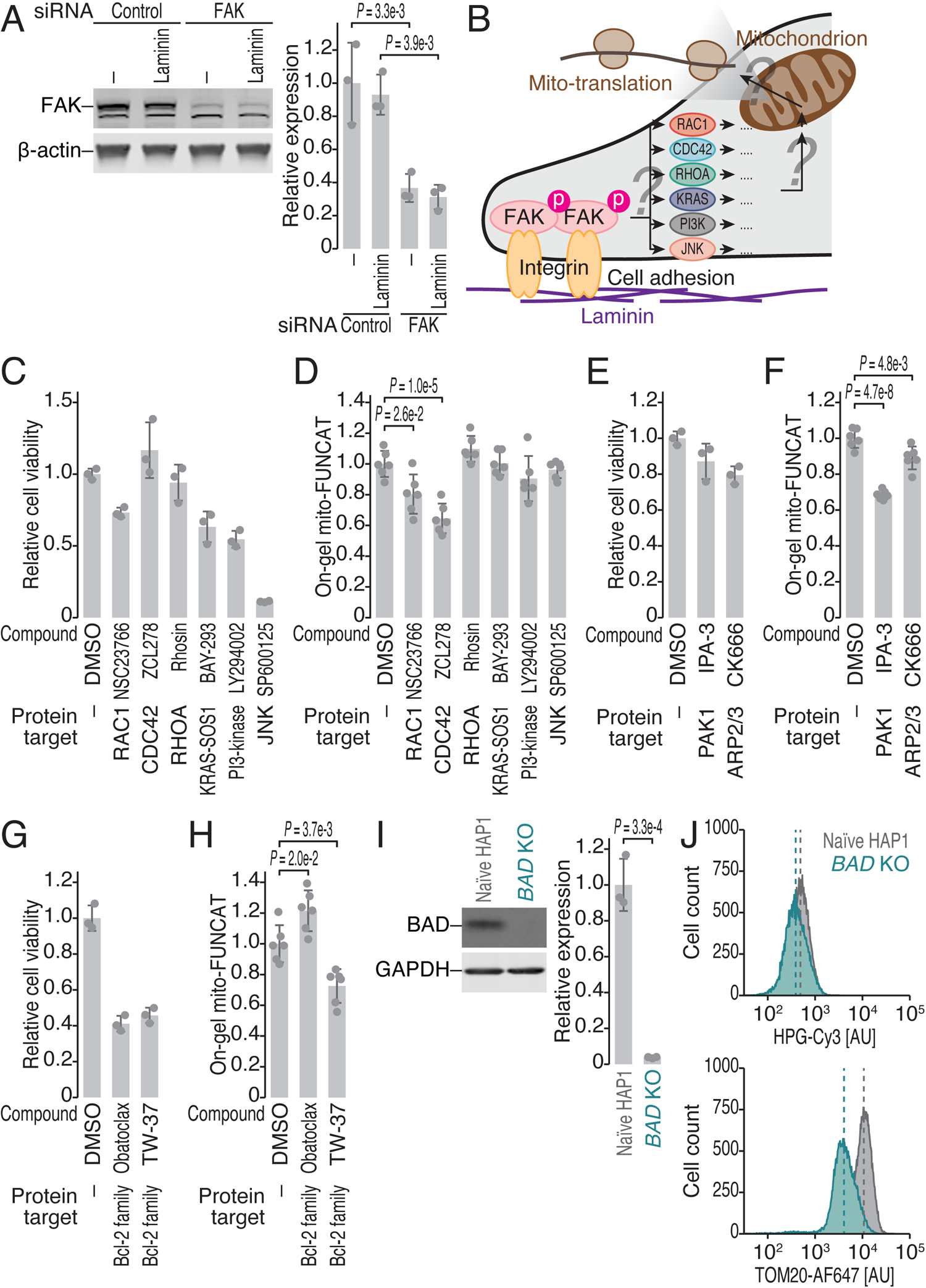
Characterization of chemical treatment for mitochondrial translation. Related to Figure 4. (A) Western blotting of FAK protein after siRNA-mediated knockdown. β-Actin was used as the loading control. The quantified relative amount of FAK is shown. (B) Schematic representation of the downstream signal transduction pathway from FAK to mitochondrial translation. (C, E, and G) Relative cell viability under treatment with the indicated compounds. The reduction in cell viability under compound treatment showed that the corresponding target proteins were efficiently blocked. (D, F, and H) On-gel FUNCAT experiments to monitor mitochondrial translation in the absence of laminin precoating under treatment with the indicated compounds. We note that the DMSO data in F are the same as the data for DMSO without laminin precoating in Figure 4D. (I) Western blotting of BAD protein in KO HAP1 cells. GAPDH was used as the loading control. The quantification of the relative amount of BAD is shown. (J) Representative distribution of Cy3-conjugated HPG signals (top) and AF647-labeled TOMM20 signals (bottom) in the indicated cell lines. The dashed line represents the mean of the distribution. For A and C-I, the data from replicates (points, n = 3 for A, C, E, G, and I; n = 6 for D, F, and H), the mean values (bars), and the s.d.s (errors) are shown. The p values were calculated by Student’s t test (two-tailed) (I) and by the Tukey‒Kramer test (two-tailed) (A, D, F, and H). For C-H, the cells were treated with the compounds at the following concentrations: NSC23766, 200 μM; ZCL278, 100 μM; Rhosin, 50 μM; BAY-293, 1 μM; LY294002, 10 μM; SP600125, 20 μM; IPA-3, 5 μM; CK666, 25 μM; obatoclax, 2 μM; and TW-37, 10 μM. The experiments were conducted in HEK293 cells unless otherwise noted.

**Figure S5.**
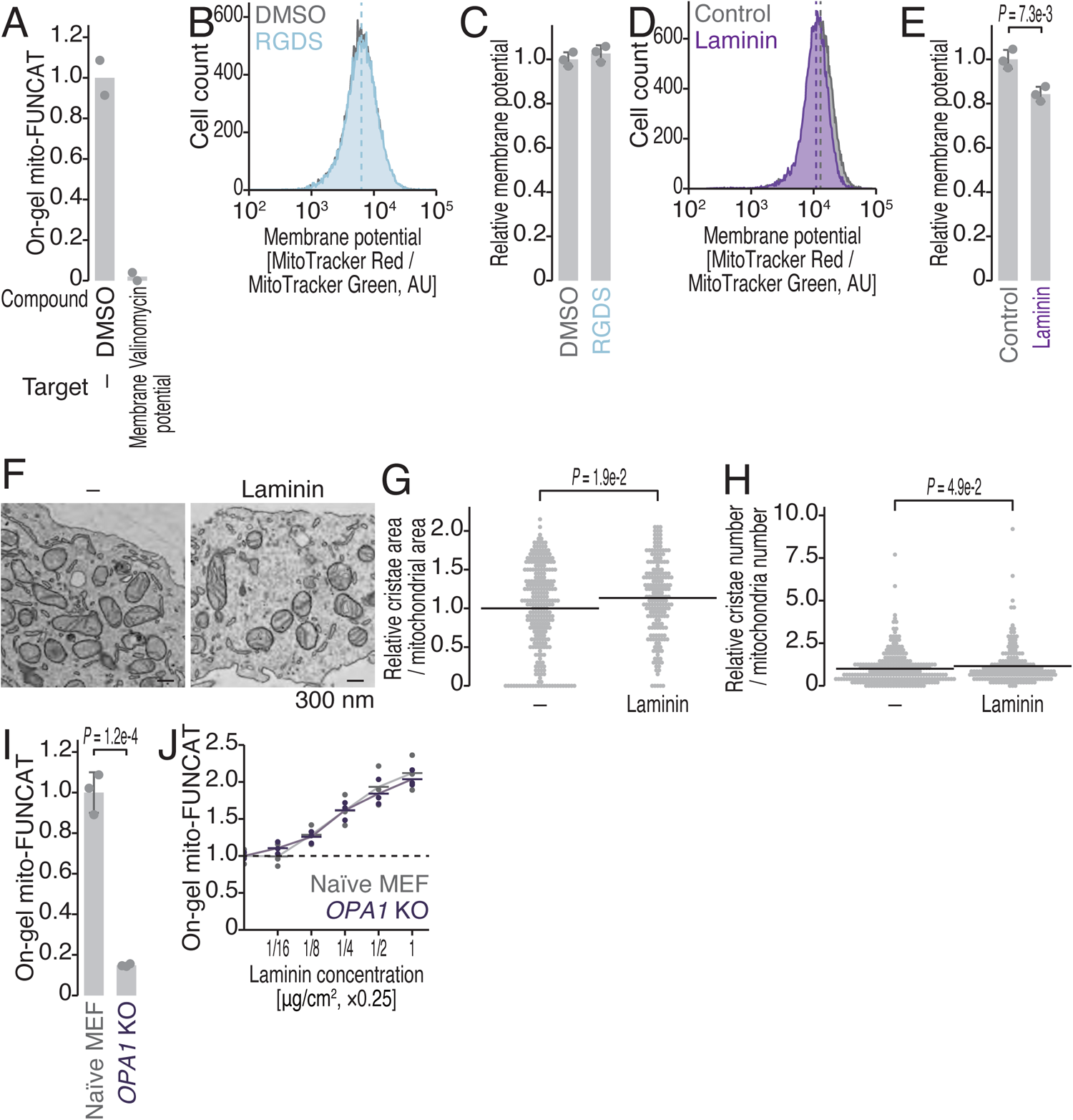
Characterization of chemical treatment for mitochondrial translation. Related to Figure 5. (A) On-gel FUNCAT experiments to monitor mitochondrial translation under treatment with the indicated compounds. (B and D) Representative distribution of MitoTracker Green-normalized MitoTracker Red values under the indicated conditions. Given that MitoTracker Red staining depends on the membrane potential, the values were normalized to MitoTracker Green, which standardizes the mitochondrial volume in cells, representing relative membrane potential. The dashed line represents the mean of the distribution. HEK293 cells were treated with RGDS peptide for 24 h. AU, arbitrary unit. (C and E) Quantification of the relative membrane potential shown in B and D. (F) Representative electron microscopy image of mitochondria under the indicated conditions. Scale bar, 300 nm. (G and H) Relative cristae area and relative cristae numbers per mitochondrion in cells precoated with laminin. The data were normalized to those of the control condition. (I and J) On-gel FUNCAT experiments to monitor mitochondrial translation in naïve and *OPA1* KO MEFs in the absence of laminin precoating (I) or with a titrated concentration of laminin for precoating (J). For A, the data from replicates (points, n = 2) and the mean values (bars) are shown. For C, E, and I, the data from replicates (points, n = 3), the mean values (bars), and the s.d.s (errors) are shown. The p values were calculated by Student’s t test (two-tailed) (E and I). For G and H, the p values were calculated by the Mann‒Whitney *U* test. For J, the data from three replicates (points) and the mean values (bars) are shown. For A, B, and C, the cells were treated with the compounds at the following concentrations: valinomycin, 2 μM; RGDS peptide, 10 μg/ml. The experiments were conducted in HEK293 cells unless otherwise noted.

**Figure S6.**
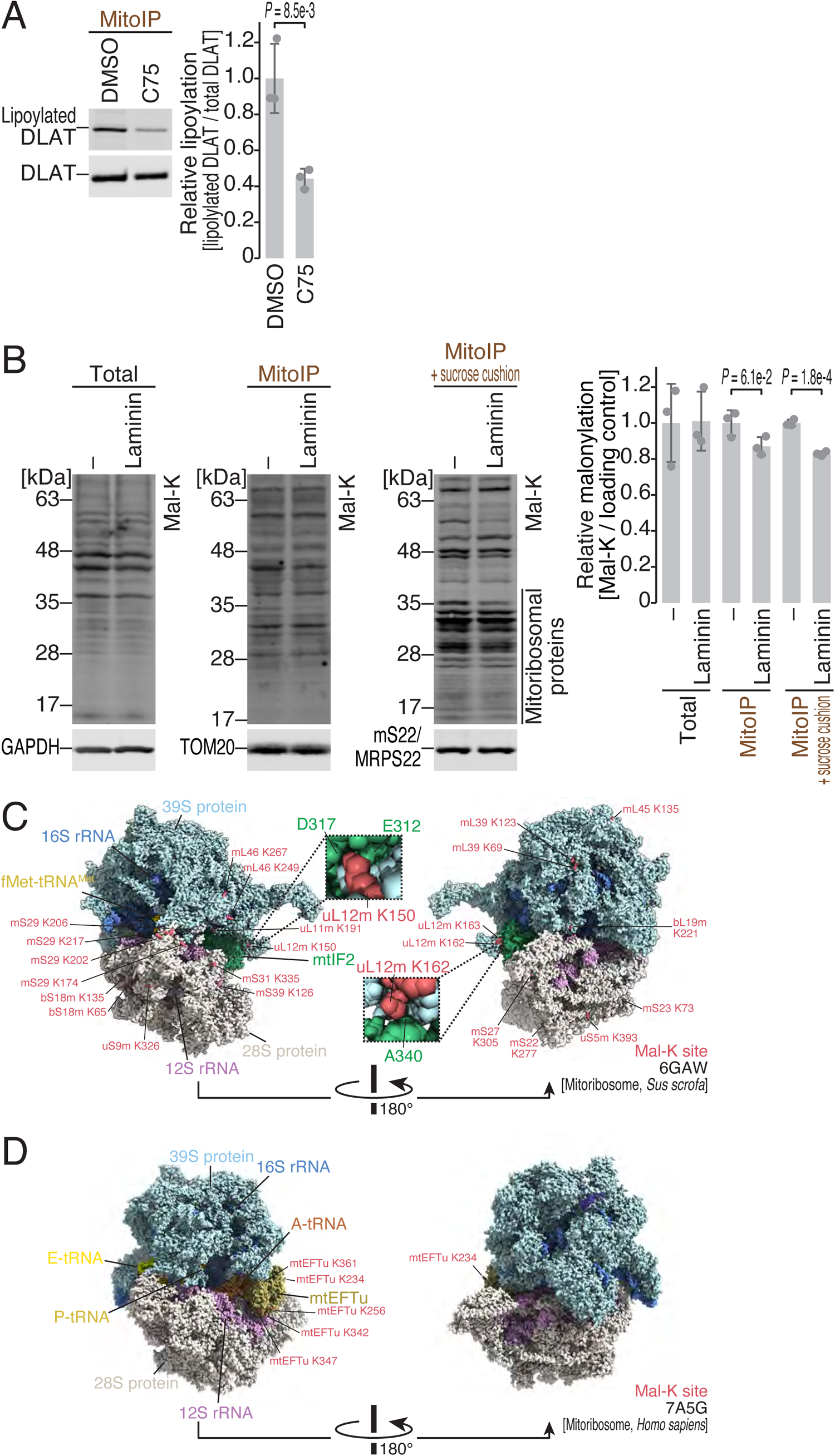
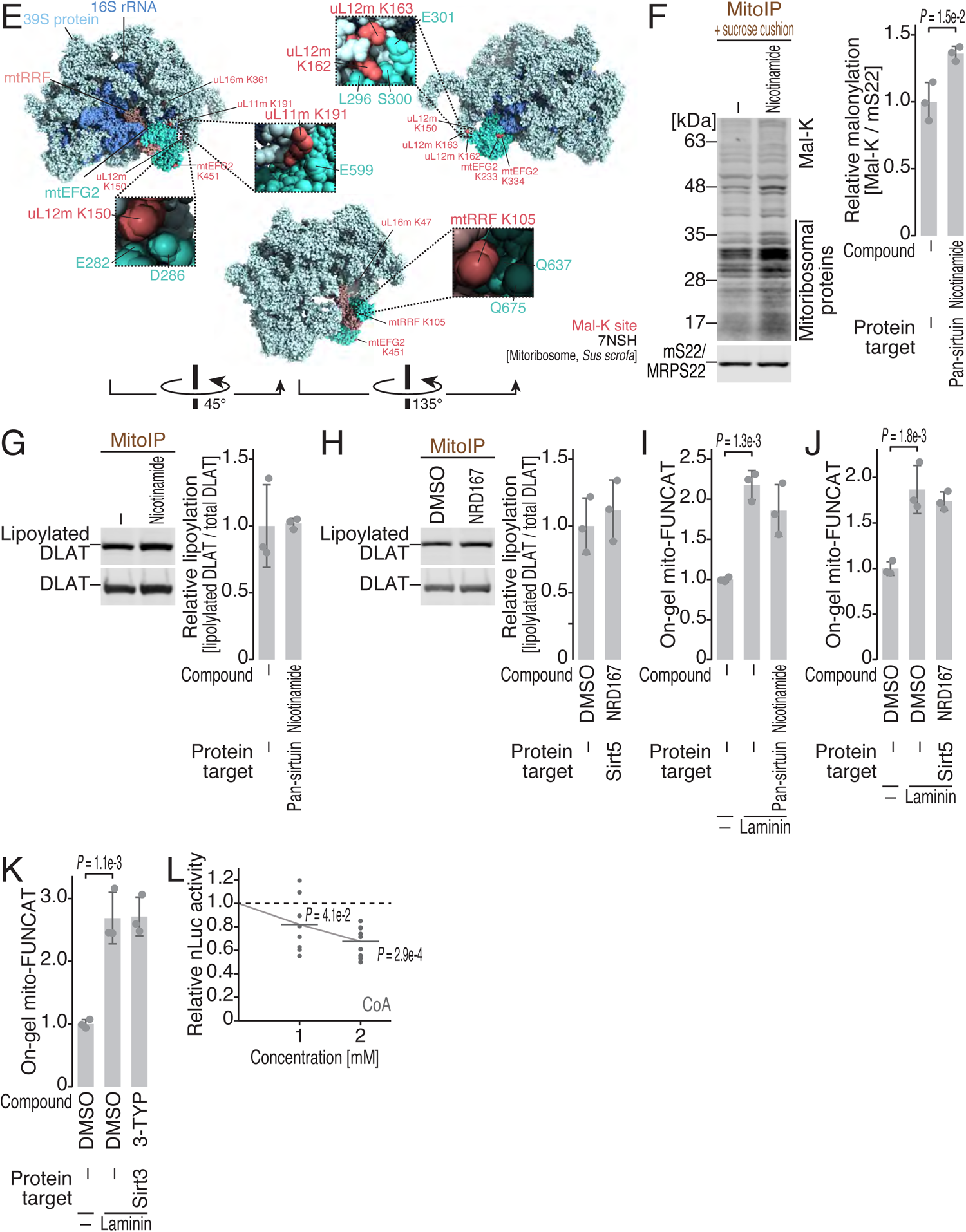
Characterization of mtFAS-coupled mitochondrial translation activation. Related to Figure 5. (A, G, and H) Western blotting of DLAT and lipoylated DLAT proteins under the indicated conditions. The quantified relative amount of lipoylated DLAT is shown. (B) Western blotting of proteins with malonylated lysine (Mal-K) in total lysate and the mitochondrial immunoprecipitation (MitoIP) fraction with an anti-TOM22 antibody with or without sucrose cushion ultracentrifugation. GAPDH, TOM20, and MRPS22 (mS22) were used as loading controls. The cells were cultured under the indicated conditions. The quantified relative amount of Mal-K signal is shown. (C-E) Malonylated lysine residues (Mal-K) found in the compendium of protein lysine modifications (CPLM) database ^68^ were mapped on the mitoribosome complex with mtIF2 (6GAW, C) ^107^, mtEFtu (7A5G, D) ^108^, and mtRRF (7NSH, E) ^109^, using UCSF ChimeraX ^106^. (F) Western blotting of proteins with malonylated lysine (Mal-K) in the mitochondrial IP (MitoIP) fraction with an anti-TOM22 antibody and then the pellet of the sucrose cushion ultracentrifugation fraction. MRPS22 (mS22) was used as the loading control. The cells were cultured under the indicated conditions. The quantified relative amount of Mal-K signal is shown. (I-K) On-gel FUNCAT experiments to monitor mitochondrial translation under treatment with the indicated compounds with or without laminin precoating. (L) *In vitro* translation of mammalian mitochondria reconstituted with recombinant factors. The effect of CoA is shown. For A, B, and F-K, the data from replicates (points, n = 3), the mean values (bars), and the s.d.s (errors) are shown. The p values were calculated by Student’s t test (two-tailed) (A-B and F) and by the Tukey‒Kramer test (two-tailed) (I-K). For L, the data from nine replicates (points) and the mean values (bars) are shown. The p values were calculated by the Tukey‒Kramer test (two-tailed). For A and F-K, the cells were treated with the compounds at the following concentrations: TW-37, 10 μM; C75, 50 μM; nicotinamide, 10 mM; NRD167, 10 μM; and 3-TYP, 10 µM. The experiments in A, B, and F-K were conducted in HEK293 cells.

**Figure S7.**
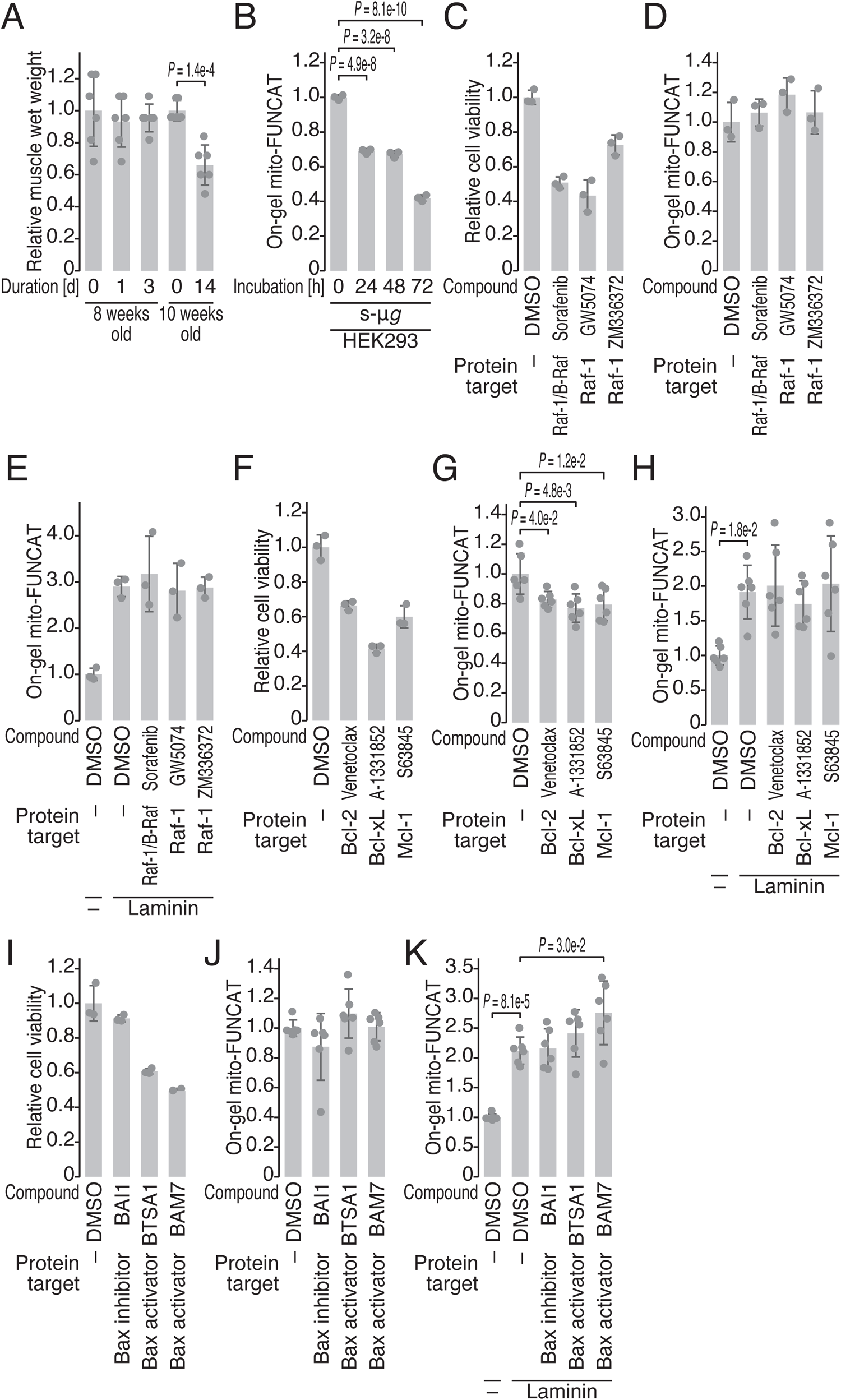
Characterization of mice under minimal mechanical stress model. Related to Figure 6. (A) Relative soleus muscle wet weight under the indicated conditions. The data were normalized to the mean value of age-matched mice without unloading treatment. The right and left knees served as different samples. (B) On-gel FUNCAT experiments to monitor mitochondrial translationover the course of simulated microgravity culture. (C, F, and I) Relative cell viability under treatment with the indicated compounds. The reduction in cell viability by compound treatment showed that the corresponding target proteins were efficiently blocked. (D, E, G, H, J, and K) On-gel FUNCAT experiments to monitor mitochondrial translation under treatment with the indicated compounds. We note that the DMSO samples in D, G, and J were the same data for DMSO without laminin precoating in E, H, and K, respectively. The data from replicates (points, n = 3 for B, C, D, E, F, and I; n = 6 for A, G, H, J, and K), the mean values (bars), and s.d.s (errors) are shown. The p values were calculated by Student’s t test (two-tailed) (A) and by the Tukey‒Kramer test (two-tailed) (B, G, H, and K). For C-K, the ells were treated with the compounds at the following concentrations: sorafenib, 1 μM; GW5074, 5 μM; ZM336372, 10 μM; venetoclax, 10 μM; A-1331852, 10 μM; S63845, 5 μM; 10 μM; BAI1, 1 μM; BTSA1, 20 μM; and BAM7, 20 μM. The experiments except for A were conducted in HEK293 cells.

**Table S1.**
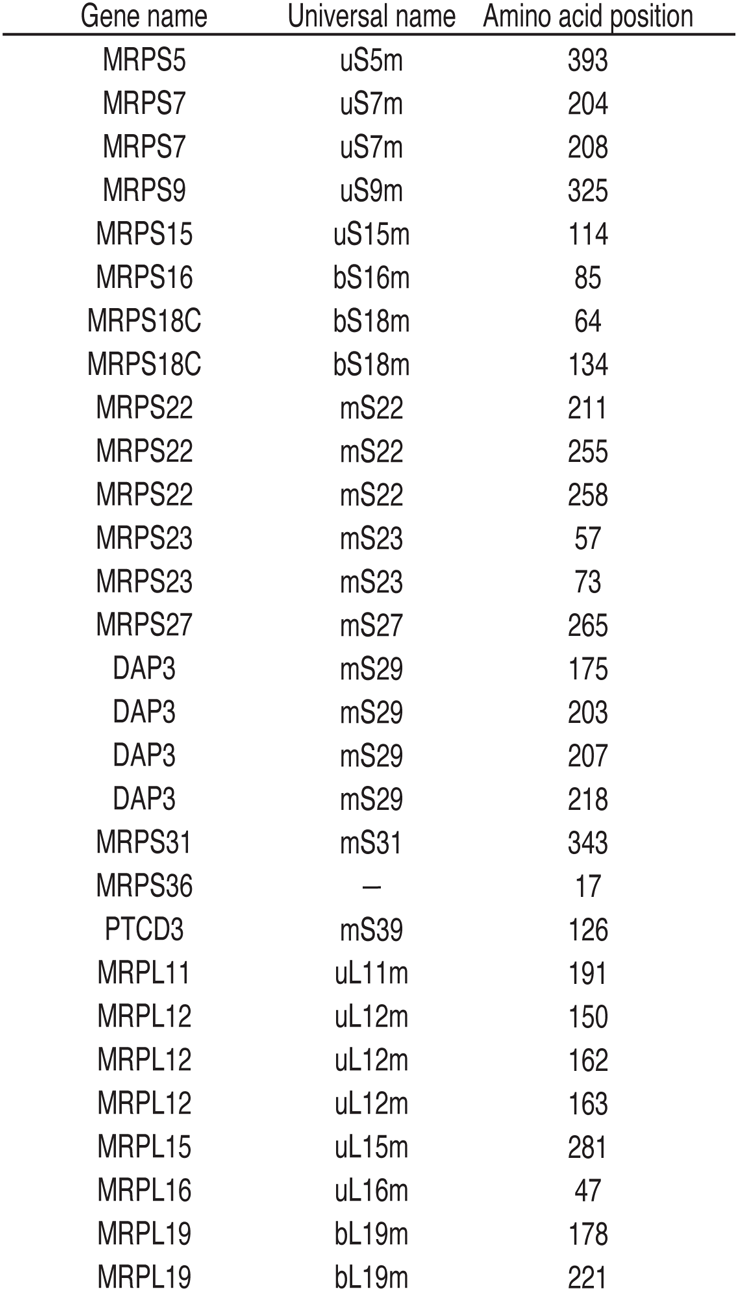
Reported Mal-K sites in the mitochondrial translation machinery. Related to Figure 5. Malonylated lysines found in the CPLM database ^68^, gene names, and universal names for ribosome proteins are listed.

**Table S2.**
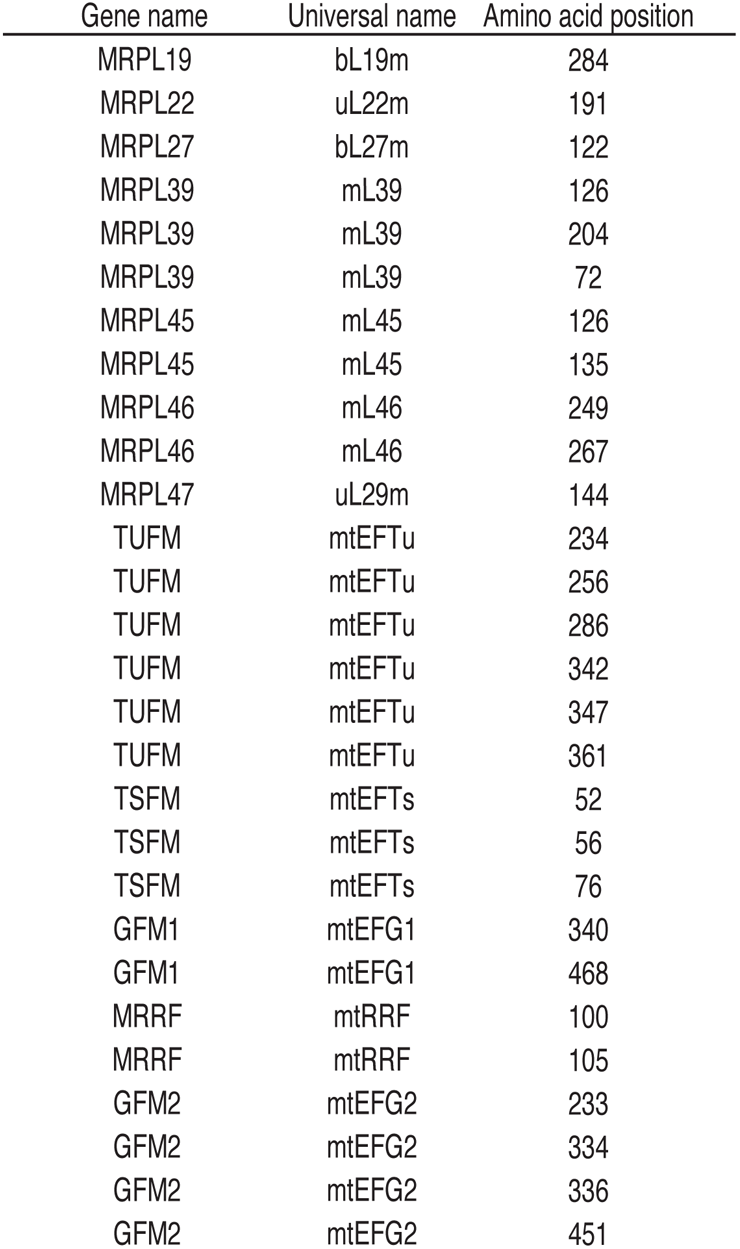
A-site offsets for cytosolic and mitochondrial ribosome footprints. Related to Figures 1, 2, 3, 5, and 6. A-site offsets for cytoribosome footprints and mitoribosome footprints along the footprint length in ribosome profiling data obtained in this study are listed.

